# A-type lamins are critical for the recruitment of RPA and RAD51 to stalled replication forks to maintain fork stability

**DOI:** 10.1101/2021.06.22.449466

**Authors:** Simona Graziano, Nuria Coll-Bonfill, Barbara Teodoro-Castro, Sahiti Kuppa, Jessica Jackson, Elena Shashkova, Urvashi Mahajan, Alessandro Vindigni, Edwin Antony, Susana Gonzalo

**Affiliations:** Edward A. Doisy Department of Biochemistry and Molecular Biology, St Louis University School of Medicine, St Louis, MO 63104; Department of Internal Medicine, Washington University School of Medicine, St Louis, MO 63106; Novo Nordisk Foundation Center for Protein Research, University of Copenhagen, Denmark

**Author notes:** Equal contribution (Simona Graziano, Nuria Coll-Bonfill). To whom correspondence should be addressed. Tel: []; Fax: []; Email: [ ].

## Abstract

Lamins provide a nuclear scaffold for compartmentalization of genome function that is important for genome integrity. The mechanisms whereby lamins regulate genome stability remain poorly understood. Here, we demonstrate a crucial role for A-type lamins preserving the integrity of the replication fork (RF) during replication stress (RS). We find that lamins bind to nascent DNA strands, especially during RS, and ensure the recruitment of fork protective factors RPA and RAD51. These ssDNA-binding proteins, considered the first and second responders to RS respectively, play crucial roles in the stabilization, remodeling and repair of the stalled fork to ensure proper restart and genome stability. Reduced recruitment of RPA and RAD51 upon lamins depletion elicits replication fork instability (RFI) depicted by MRE11 nuclease-mediated degradation of nascent DNA, RS-induced accumulation of DNA damage, and increased sensitivity to replication inhibitors. Importantly, in contrast to cells deficient in various homology recombination repair proteins, the RFI phenotype of lamins-depleted cells is not linked to RF reversal. This suggests that the point of entry of nucleases is not the reversed fork, but regions of ssDNA generated during RS that are not protected by RPA and RAD51. Consistently, RFI in lamins-depleted cells is rescued by forced elevation of the heterotrimeric RPA complex or RAD51. These data unveil a clear involvement of structural nuclear proteins in the protection of ssDNA from the action of nucleases during RS by warranting proper recruitment of ssDNA binding proteins RPA and RAD51 to stalled RFs. In support of this model, we show physical interaction between RPA and lamins. Our study also suggests that RS is a major source of genomic instability in laminopathies and in lamins-depleted tumors.

## INTRODUCTION

The spatial organization of the genome, orchestrated by structural nuclear proteins such as A-type and B-type lamins, is important for genome function and integrity. Lamins maintain nuclear architecture, a proper response to mechanical stress, chromatin organization, and genome stability (1,2). Lamins dysfunction is associated with degenerative disorders, premature aging, and cancer, suggesting that these proteins operate as “genome caretakers”. In particular, A-type lamins (lamin-A/C, herein lamins) dysfunction causes alterations in telomere biology and DNA repair mechanisms (3–9). Although DNA replication is the major source of spontaneous DNA damage in dividing cells (10,11), the role of lamins in replication has been minimally explored. Early studies showed colocalization of lamins with PCNA (12–14) and a direct association with DNA polymerases Pol α, δ, and e (15), suggesting a role for lamins in replication. Other studies showed that lamins are required for restart of stalled replication forks (RFs) (16), and that mutant forms of lamin A sequester PCNA away from the RF (17–19). Despite these findings, our mechanistic understanding of lamins function in replication is limited.

During replication, the fork encounters many challenges, including DNA lesions and secondary structures in DNA that cause RF stalling and replication stress (RS) (11). Upon fork stalling, there is uncoupling of DNA polymerases from the MCM helicase complex, which generates regions of ssDNA that are rapidly coated by the heterotrimeric RPA complex (RPA70, RPA32 and RPA14 subunits) (20). The ssDNA-RPA complex plays a key role during RS, activating the ATR/Chk1-dependent S-phase checkpoint, which phosphorylates among other factors, chromatin-bound RPA32 on multiple sites including S33 (21–23). This complex acts as a platform for recruitment of factors such as BRCA1/2, FANCD2, PALB2, and ultimately RAD51, which replaces RPA on ssDNA and forms a filament that stabilizes and remodels the stalled fork (24,25). These proteins, classically associated with homologous recombination (HR) repair, protect the stalled fork from degradation by nucleases such as MRE11 (26–28). RPA and RAD51 have also been shown to facilitate fork reversal, a key protective mechanism that involves the annealing of two newly synthesized strands to form a chicken-foot structure (29) that allows forks with persistent ssDNA gaps to reverse their course and resume replication without chromosomal breakage (30,31). RPA promotes SMARCAL1 binding to RF, which is required for remodeling forks into reversed structures (32). RAD51 is thought to have two functions in the reversal of stalled RF: (1) promote the initial step of fork regression/reversal in a BRCA-independent manner; and (2) stabilize the already formed reversed fork from degradation in a BRCA-dependent manner (29,33,34). In addition, RAD51 mediates the restart of stalled RFs and repair of DNA damage (35). Interestingly, the protective function of RAD51 from MRE11-mediated degradation and the remodeling of stalled forks does not seem to require its enzymatic recombinase activity (36). In the case of RPA, non-canonical roles include functioning as a sensor of R-loops (RNA:DNA hybrids with ssDNA), which represent a barrier to replication fork progression. RPA was shown to stimulate the activity of RNaseH1 suppressing R loops (37). Moreover, RPA recruitment to sites of LINE-1 integration via binding to Poly(ADP)-Ribose Polymerase 2 (PARP2) facilitates retrotransposition (38). These studies showcase the versatility of RPA and RAD51 in the maintenance of genomic stability, and their crucial tasks in RF protection during RS. The nuclear pools of RAD51 and RPA are finite and rate limiting for ssDNA protection at stalled forks. Excessive accumulation of ssDNA in cells, for example by ATR inhibition, TREX1 deficiency, or toxigenic bacterial infection, overwhelms the RPA and/or RAD51 response to DNA damage, leading to exhaustion of RPA and RAD51 nuclear pools and an increase in DNA breakage that stems from RS (39–41).

Given the roles of structural proteins “A-type lamins” in telomere biology and DNA repair (3-9), we determined if lamins play a role in DNA replication. Here, we show for the first time that lamins are essential for the efficient recruitment of RPA and RAD51 to stalled RFs during RS, and that their reduced recruitment upon lamins loss causes RFI to the same extent as HR-deficient cells. Lamins physically interact with RPA and likely act to recruit both RPA and RAD51 to RFs. Accordingly, lamins depletion elicits nuclease-mediated degradation of stalled forks, RS-induced genomic instability, and increased sensitivity to drugs that inhibit replication. Overall, our studies indicate that reduced expression of lamins, which is associated with poor prognosis in many cancers, elicits phenotypes of genomic instability due to degradation of ssDNA during replication.

## MATERIAL AND METHODS

### Cell Culture

HEK293 cells and primary MEFs from *Lmna*^+/+^ and *Lmna*^−/−^ were immortalized with SV40LT. All these cells and cancer cells -U2OS, MCF-7 and MDA-MB-231-were cultured in DMEM supplemented with 10% FBS+1% antibiotics/antimycotics.

### Constructs and Viral Transduction

Retroviral and lentiviral transductions were performed as previously described (5). Viral envelope and packaging plasmids were gifts from Sheila Stewart (WUSM), shRNAs targeting Lamin-A/C and BRCA2 were purchased from Sigma Aldrich, lamin-A and progerin expressing plasmids were a gift from Brian Kennedy (Buck Institute, CA and National University Singapore), and RAD51 construct from Simon Powell (Memorial Sloan Kettering). Constructs for expression of the heterotrimeric RPA complex (super-RPA) were provided by Luis Toledo (University of Copenhagen).

### RNA interference

Transient transfection of siRNAs was performed using Lipofectamine RNAiMax transfection reagent (Life Technologies). SMARTpool siRNA Dharmacon were used to deplete BRCA2 (L-003462-00, 50nM, 48h), MRE11 (L-009271-00, 50nM, 48h), SMARCAL1 (D-013058-02-0002, 50nM, 48h), while RAD51 was depleted by employing siRNA Ambion (4390827, 50nM, 48h) as described (28). In each experiment, silencer select negative siRNA (4390843, Ambion) was used as transfection control.

### DNA Fiber Assays

Fiber assays were performed as previously described (42), with some modifications. Briefly, asynchronous cells were labeled with thymidine analogs: 20 μM iododeoxyuridine (ldU) followed by 200 μM chlorodeoxyuridine (CIdU). The specific labeling scheme and treatments for each experiment are shown in figure legends. Cells were collected by trypsinization, washed and resuspended in 100 μl of PBS. Then, 2 μl cell suspension was dropped on a polarized slide (Denville Ultraclear) and cell lysis was performed in situ by adding 8 μl lysis buffer (200 mM Tris-HCl pH 7.5; 50 mM EDTA; 0.5% SDS). Stretching of high-molecular weight DNA was achieved by tilting the slides at 15-45°. The resulting DNA spreads were air dried and fixed for 5 min in 3:1 Methanol:Acetic acid and refrigerated overnight. For immunostaining, stretched DNA fibers were denatured with 2.5 N HCl for 60 min, washed 3 × 5 min in PBS, then blocked with 5% BSA in PBS for 30 min at 37°C. Rat anti-CldU/BrdU (Abcam, ab6326) (1:100), chicken anti-rat Alexa 488 (Invitrogen, A21470) (1:100), mouse anti-IdU/BrdU (BD Biosciences, 347580) (1:20) and goat anti-mouse IgG1 Alexa 547 (Invitrogen, A21123) (1:100) antibodies were used to reveal CldU- and IdU-labeled tracts, respectively. Labeled tracts were visualized under a Leica SP5X confocal microscope using 63x oil objective lens (NA 1.40, with LAS AF software), and tract lengths were measured using ImageJ. A total of 100-300 fibers were analyzed for each condition in each experiment. All analyses were performed blinding the samples. Statistical analysis of the tract length was performed using GraphPad Prism.

#### Labeling schemes and treatments

To monitor DNA end resection, cells are labeled with thymidine analogs for equal amounts of time: 20 min IdU followed by 20 min CldU and then exposed for 3-5 hours to either 4 mM HU, 50 μM Mirin, 4 mM HU + 50 μM Mirin, 200 nM Aphidicolin, 100 nM Camptothecin, or vehicle. To exclude that end-resection was an artifact of the used labeling scheme, progressive forks have been first labeled for 20 min with IdU and then exposed for 2 hours to 4 mM HU+CldU or 4 mM HU + 50 μM Mirin+CldU.

### Analysis of cell cycle profile

Asynchronous and S-phase synchronized cells pulse-labeled with the thymidine analog EdU for 20 min were trypsinized, washed once with PBS, and then suspended in 500 μl of PBS prior to the dropwise addition of 500 μl of formaldehyde 7.4% and incubated at room temperature for 10 min. Fixed cells were washed once with PBS, blocked in 1% BSA/PBS for 10 min, pelleted at 2000 rpm, and permeabilized in 0.5% saponin/1% BSA/PBS for 30 min at room temperature. Cells were then pelleted at 2000 rpm, washed with 1% BSA/PBS once, pelleted at 2000 rpm and incubated in the Click-it reaction solution prepared according to manufacturer (Click-it EdU Alexa Fluor 488 Imaging Kit, life technologies #C10337) for 30 min at room temperature in the dark. Cells were washed with 1% BSA/PBS once, pelleted at 2000 rpm and incubated with DAPI staining solution for 30 min. Stained cells were analyzed for fluorescent DNA content and EdU content using the BD Biosciences LSR II flow cytometer, and the cell cycle profiles were created by the program FlowJo^®^. <u>DNA staining solution</u>: 1% BSA/PBS, 0.1 mg/ml RNase (Thermo-Scientific 12091-039), 2 μg/ml DAPI.

### Subcellular fractionation

Performed using “Subcellular protein fractionation kit for cultured cells” from Fisher Thermo Scientific according to manufacturer instructions. The day before, 5X10^6^ cells were seeded in 150 mm dishes and cultured overnight in complete medium. Cells were left asynchronous (control) or synchronized in S-phase by overnight exposure to 1 μM aphidicolin (APH). The next day synchronized cells were released for 2 hours in complete media and either exposed to 4 mM HU or vehicle for 2 hours prior to subcellular fractionation.

### Immunoblotting

Immunoblotting was carried out by lysing cells in UREA buffer (8 M Urea, 40 mM Tris pH 7.5, 1% NP40), for 10 min on ice. Lysates were centrifuged at 14000 rpm for 12 min at 4°C, and the DNA pellet removed from the sample. 80 μg of total protein was separated by SDS-PAGE on a 4-15% Criterion TGX Gel (Bio-Rad) and transferred to a nitrocellulose membrane using the Trans-Blot Turbo system (Bio-Rad). Membranes were blocked using 5% BSA in PBS+0.1% Tween-20 for 1 hour at room temperature, then incubated overnight at 4°C with the appropriate antibody diluted in blocking solution. Membranes were washed 3X using PBS+0.1% Tween-20 after both primary and secondary antibody incubations. Membranes were developed using Immobilon Western Chemiluminescent HRP Substrate (Merck Millipore).

Antibodies used: BRCA1 (Calbiochem OP93), RAD51 (H-92: sc-8349 SCBT; AB3756 Millipore), 53BP1 (H-300: sc-22760 SCBT), Vinculin (N-19: sc-7649 SCBT), γH2AX (cs-2577, Cell Signaling), Lamin-A/C (H-110: sc-20681 SCBT), RPA (Ab-2: Calbiochem #NA18), P-RPA32 on S4/S8 (Bethyl A300-245A), P-RPA on S33 (Bethyl A300-246A), BRCA2 (Abcam Ab-1: OP95 Ab90451), MRE11 (Novus Biological NB100-142), PCNA (PC-10: sc-56 SCBT), CHK1 (sc8408 SCBT), P-CHK1 (Cell Signaling 2348s), Histone 3 (Ab1791, AbCam), SMARCAL1 (sc376377 SCBT).

### Immunofluorescence

Approximately 5X10^5^ cells were treated with HU (4mM for 4h) or vehicle. After washing, cells were incubated 10 min on ice with an extraction buffer (25mM HEPEs pH 7.4, 50 mM NaCl, 1 mM EDTA, 3 mM MgCl2, 300 mM Sucrose 0.5% TX-100 in ddH20). After 3 washes in 1X PBS, cells were fixed in 4% formaldehyde in PBS for 10 min, permeabilizated in 0.5% TX-100 30 min at room temperature and blocked 1 h at 37°C in 1% BSA/PBS. Incubation with primary antibody was performed ON at 4°C and secondary antibody for 1h at 37°C, both in a humid chamber. After washes in PBS, cells were counterstained with DAPI in Vectashield. Microscopy and photo capture were performed on a Leica DM5000 B microscope using 40× or 63× oil objective lenses with a Leica DFC350FX digital camera and the Leica Application Suite. Antibody used: P-RPA on S33 (Bethyl A300-246A).

### Isolation of proteins at nascent DNA (iPOND)

iPOND was performed as originally described (Sirbu et al., 2012) with minor modifications. HEK293T cells were pulse-labeled with 10 μM EdU (Life Technologies) for 15 min and left untreated or treated with 4 mM HU for 2 hours. For the thymidine (Thy) chase experiments, cells were washed 3X with complete medium and incubated for 60 min in medium supplemented with 10 μM (Thy) (Sigma-Aldrich). Protein-DNA crosslinking was performed in 1% formaldehyde for 15 min at RT, quenched with 0.125 M glycine for 5 min, and washed 3X with PBS. Cells were then permeabilized with 0.25% Triton X-100/PBS for 30 minutes, washed once with 0.5% BSA/PBS and once with PBS, and incubated in click reaction buffer (10 mM sodium-l-ascorbate, 20 μM biotin azide [Life Technologies], and 2 mM CuSO4) for 2h at room temperature. In the “no-click” control sample DMSO was used in place of biotin azide in the click reaction buffer. Cells were washed once with 0.5% BSA/PBS and once with PBS prior to be resuspended in lysis buffer (50 mM Tris-HCl, pH 8.0, and 1% SDS) supplemented with protease/phosphatase inhibitor (Cell Signaling), and chromatin was solubilized by sonication in a Bioruptor (Diagenode) at 4°C at the highest setting for 30-45 cycles (30sec on and 30sec off). Samples were centrifuged for 10 min at 5000Xg and supernatants were diluted 1:1 with PBS (vol/vol) containing protease/phosphatase inhibitors and incubated overnight at +4°C with streptavidin-agarose beads (EMD Millipore). Beads were washed twice with lysis buffer, once with 1 M NaCl, twice with lysis buffer and captured proteins were eluted by boiling beads in 2X NuPAGE LDS Sample Buffer (Life Technologies) containing 200 mM DTT for 40 min at 95°C. Proteins were resolved by electrophoresis using BioRad 4–12% Bis-Tris gels and detected by Western blotting.

### Metaphase spreads

Cells were cultured in 10 mm dishes to 70% confluence and asynchronous cells were exposed to DMSO, 4 mM HU, 50 μM Mirin, or 4 mM HU + 50 μM Mirin for 5 hours. Cells were allowed to complete S-phase in complete medium supplemented with 10 μM Colcemid (Sigma #D1925) in order to arrest them in metaphase. After collecting the culture media, cells were washed with 1X PBS (which was also collected), and the trypsinized cells were collected. All the fractions were combined and centrifuged to pellet the cells. Supernatant was then aspirated to leave ~2 ml of media + cell pellet, which was resuspended by gentle flicking. 9 ml of pre-heated (37°C) hypotonic buffer (0.56% KCl) was added then to the cell suspension, which was incubated in a 37°C water bath to allow hypotonic swelling of the cells. A small amount (~500 μl) of fixing solution (3:1 methanol:acetic acid) was added and the cells were pelleted at 4°C by centrifugation. Cells were kept on ice from this point onwards. After aspirating the supernatant to leave ~2 ml, the cells were resuspended by gentle flicking and 9 ml fixing solution was added dropwise. The suspension was centrifuged once more to pellet the cells and fixing solution added in a similar manner. This mixture was stored at −20°C overnight. Cells were then pelleted, resuspended in ~500 μl of fresh fixing solution, dropped onto microscope slides, and allowed to air-dry at room temperature overnight. Metaphase spreads were stained by dipping the microscope slides into a Coplin jar containing a solution of Giemsa (Sigma #GS500) diluted in milliQ water 1:20, for 20 min. The excess staining was washed off with tap water and slides air-dried at room temperature. Metaphase spreads were visualized under a Leica DM5000 B microscope using 100X oil objective lens (NA 1.3) with a Leica DFC350FX digital camera and the Leica Application Suite (Version 4.1.0).

### Comet Assay

Neutral comet assays were performed using CometSlide assay kits (Trevigen). Cells were treated with either vehicle (DMSO), 4 mM HU, 50 μM Mirin, or HU+Mirin for 5 hours. Cells were embedded in agarose, lysed and subjected to neutral electrophoresis. Before image analysis, cells were stained with ethidium bromide and visualized under a fluorescence Leica DM5000 B microscope using 20X objective lens (NA 0.5) with a Leica DFC350FX digital camera and the Leica Application Suite (Version 4.1.0). Single-cell electrophoresis results in a comet-shaped distribution of DNA. The comet head contains high molecular weight and intact DNA, and the tails contains the leading ends of migrating fragments. Olive comet moment was calculated by multiplying the percentage of DNA in the tail by the displacement between the means of the head and tail distributions, as described (43). We utilized the program CometScore™ version 1.5 (TriTek) to calculate Olive Comet Moment. A total of 60-100 comets were analyzed per sample in each experiment.

### XTT assay

Cell viability monitored using “Cell proliferation kit II (XTT)” from Sigma according to manufacturer instructions. Cells were plated in 96-well dishes at low confluency and cultured for 6 days in complete medium supplemented with different concentrations of the replication inhibitor HU (50, 100, 200, 400 μM) or the PARP inhibitor Olaparib (1, 10, 50, 100 μM). Drugs were freshly replaced every other day.

### EM analysis

The procedure was performed as described (44). Lamins-proficient and -deficient cells were treated with HU and Mirin for the indicated times. In vivo psoralen cross-linking of the DNA was achieved by a repetitive exposure of living cells to 4,5’,8-trimethylpsoralen (10 μg/ml final concentration) followed by irradiation pulses with UV 365-nm monochromatic light (UV Stratalinker 1800; Agilent Technologies). The cells were then lysed with cell lysis buffer (buffer C1: 1.28 M sucrose, 40 mM Tris-Cl, pH 7.5, 20 mM MgCl_2_, and 4% Triton X-100; QIAGEN) and then digested by digestion buffer (QIAGEN buffer G2: 800 mM guanidine-HCl, 30 mM Tris-HCl, pH 8.0, 30 mM EDTA, pH 8.0, 5% Tween 20, and 0.5% Triton X-100) and 1 mg/ml proteinase K at 50°C for 2 h. Chloroform/Isoamyl alcohol (24:1) was used to collect DNA via phase separation (centrifugation at 8,000 rpm for 20 min) followed by DNA precipitation by adding 0.7× volume of isopropanol. The DNA was then washed with 70% ethanol, air dried, and resuspended in 200 μl TE (Tris-EDTA) buffer. 100 U restriction enzyme PvuII high-fidelity was used for 12 μg mammalian genomic DNA digestion (4-5-h incubation). Poly-Prep chromatography columns were used for RI enrichment. Benzoylated naphthoylated DEAE-cellulose granules were resuspended in 10 mM Tris-HCl, pH 8.0, and 300 mM NaCl to a final concentration of 0.1 g/ml. The columns were washed and equilibrated with 10 mM Tris-HCl, pH 8.0, and 1 M NaCl and 10 mM Tris-HCl, pH 8.0, and 300 mM NaCl, respectively. The sample DNA was then loaded and incubated for 0.5 h. After washing the columns (10 mM Tris-HCl, pH 8.0, and 1 M NaCl), the DNA was eluted in caffeine solution (10 mM Tris-HCl, pH 8.0, 1 M NaCl, and 1.8% [wt/vol] caffeine) for 10 min followed by sample collection. DNA is then purified and concentrated, using an Amicon size-exclusion column and resuspended in TE. With DNA spreading by the “BAC method,” the DNA was loaded on carbon-coated 400-mesh copper grids. The DNA was then coated with platinum by platinum-carbon rotary shadowing (High Vacuum Evaporator MED 020; Bal-Tec). Microscopy was performed with a transmission electron microscope (Tecnai G2 Spirit; FEI; LaB6 filament; high tension ≤120 kV) and acquired with a side mount charge-coupled device camera (2,600 × 4,000 pixels; Orius 1000; Gatan, Inc.). The images were processed with DigitalMicrograph Version 1.83.842 (Gatan, Inc.) and analyzed with ImageJ (National Institutes of Health).

### Statistical analysis

All data sets from each DNA fiber assay experiment have been first subjected to normality test. Most of the data set does not meet the criteria of a normal distribution. Since each data set includes more than 100 values to calculate statistical significance of the observed differences we have used: nonparametric tests, such as Wilcoxon matched-pairs signed rank test for paired samples and Mann-Whitney test for unpaired samples; one-way Anova was used for all the experiments in which more than two comparisons were performed. GraphPad Prism 6 was used for the calculations. In all cases, differences were considered statistically significant when p<0.05.

### RPA binding to Lamin

Human RPA, RPA-FAB, and RPA-32-14 proteins were purified as described (45). Lamin-C was overproduced and purified from codon-optimized *Lamin-C* (amino acid residues 1-152) engineered into pRSF-Duet plasmid (Genscript Inc.). The construct carries a tandem N-terminal Strep-tag, polyhistidine tag (6X His), and a SUMO-tag along with a SUMO protease cleavage site engineered after the tags. The plasmid was transformed into BL21-pLysS cells and a 10 ml overnight culture from a single transformant was grown in Luria Broth media. Larger 1 L cultures were grown from the starter culture and at 37°C until OD_600_ reached 0.6 and Lamin-C overproduction was induced with 0.5 mM IPTG. Induction was carried out at 37°C for 4 hours. Harvested cells were resuspended in 120 ml cell resuspension buffer (50 mM HEPES, pH 8.0, 750 mM NaCl, 5 mM βME, 1.5X protease inhibitor cocktail, 1 mM PMSF, 10% (v/v) glycerol and 10 mM imidazole). Cells were lysed using 0.4 g/ml lysozyme followed by sonication. Clarified lysates were fractionated on a Ni^2+^-NTA agarose column. Bound proteins were eluted using a step-gradient in cell resuspension buffer containing 0-500 mM imidazole. Fractions containing Lamin-C were pooled and concentrated using an Amicon spin concentrator (30 kDa molecular weight cut-off). Concentrated Lamin-C was dialyzed against Lamin-C storage buffer (50 mM HEPES, pH 8.0, 750 mM NaCl, 5 mM βME, 1.5X protease inhibitor cocktail, 10 % (v/v) glycerol). Lamin-C was flash frozen using liquid nitrogen and stored at −80°C. RPA concentration was measured spectroscopically using e_280_ = 87,210 M^-1^cm^-1^ (hRPA-WT), 42,400 M^-1^cm^-1^ (hRPA-FAB), or 24,410 M^-1^cm^-1^ (hRPA-32-14). The concentration of Lamin-C was calculated using the Bradford method with BSA as reference.

Lamin-C binding to RPA was captured using Bio-layer interferometry (BLI). Experiments were performed using a single channel BLItz instrument (Sartorius Inc.) and binding sensograms were collected in advanced kinetics mode at room temperature with shaking at 2200 rpm. Strep-tagged Lamin-C was bound onto streptavidin (SA) biosensors (Sartorious Inc.) which were pre-hydrated by incubating the tips in BLI buffer (30 mM HEPES, pH 7.8, 100 mM KCl, 5 mM MgCl_2_, 6 % v/v glycerol) for 10 mins. Binding and dissociation of RPA, RPA-FAB or RPA-32-14 to the Lamin-C-bound tip was measured. Sensograms were fit using the associated BLItz Pro software to obtain K_d_ values.

## RESULTS

### Lamins loss causes nucleolytic degradation of stalled replication forks

DNA lesions caused by endogenous and exogenous agents, secondary DNA structures, and transcribing RNA polymerases, pose a challenge for the RF. Damaged RFs stall and recruit factors that protect them from excessive nucleolytic degradation, while facilitating fork restart to preserve genome stability (11,25,31). To determine how lamins loss impacts RF progression and stability upon stalling, we performed DNA fiber assays in HEK-293T and MCF7 cells transduced with shRNAs targeting lamin-A/C (shLmna) or luciferase (shLuc) as control (**Fig 1A**). Replication events were labeled with thymidine analogs: IdU (iododeoxyuridine/red tract) followed by CldU (chlorodeoxyuridine/green tract) or viceversa (**Fig S1**), for an equal amount of time and detected by immunofluorescence (46). Progressing forks were identified by continuous IdU-CldU label and the length of red (IdU) and green (CldU) tracts measured. We find that lamins depletion does not affect RF progression, as shown by the similar length of red and green tracts (**Fig S1**), and an average CldU/IdU ratio of ~1 (**Fig 1B**). However, treatment with 4 mM hydroxyurea (HU) for three hours to slow-down/stall RFs results in shortening of the most recently synthesized DNA filament (CdU tract), and therefore decreased CldU/IdU ratio in lamins-depleted cells (**Fig 1B**). Thus, lamins depletion hinders the stability of stalled RFs. Failure to protect stalled RFs causes nucleolytic degradation, a typical phenotype displayed by cells deficient in factors that promote loading of RAD51 on ssDNA like BRCA1/2 and FANCD2, cells lacking RAD51 mediators (RAD51C, XRCC2), and cells with partial inhibition of RAD51 activity or expression (24,26,33,47). In all these contexts there is inefficient nucleation and/or stabilization of RAD51 protofilaments and DNA is exposed to the action of nucleases such as MRE11, EXO1 and DNA2 (36). Since MRE11 initiates end resection while EXO1 and DNA2 are involved in later steps of long-range degradation (48), we sought to determine whether MRE11 is responsible for fork degradation in our cells. We find that green tract shortening upon fork stalling in lamins-depleted cells is due to MRE11-mediated degradation, as treatment with Mirin, an inhibitor of MRE11 nuclease, rescues fork resection (**Fig 1B**). Consistently, MRE11 depletion by siRNA restores the average ratio CldU/IdU to ~1 in lamins-depleted cells (**Fig 1C**). Importantly, lamins loss does not affect cell cycle progression, as indicated by the normal cell cycle profile observed by flow cytometry (**Fig S2**). Similarly, the three-hour treatment with HU and Mirin does not alter the percentage of cells in S-phase. These control experiments support the validity of our cellular model and experimental conditions to study DNA replication.

**Figure 1.**
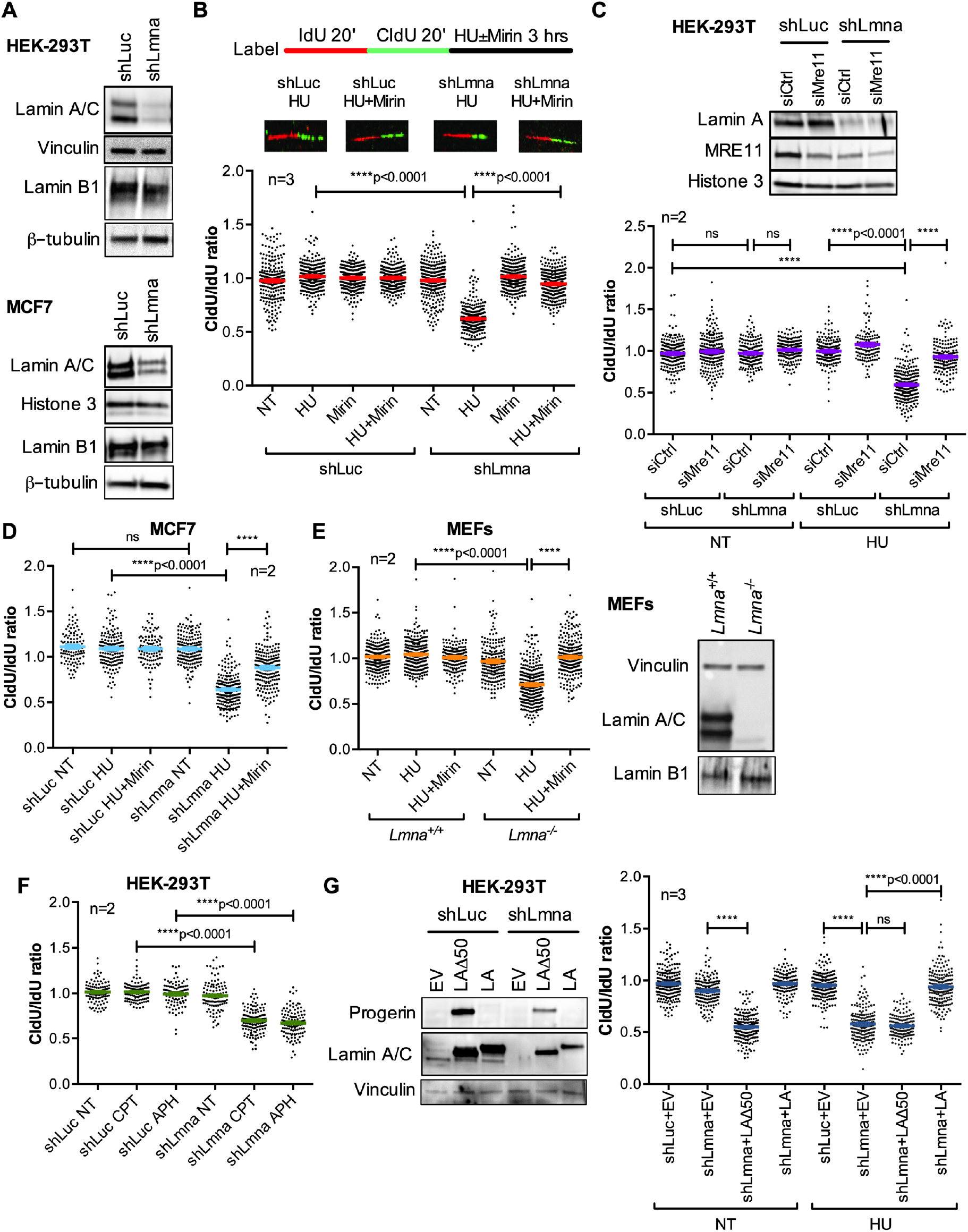
Lamins loss results in nuclease-mediated degradation of stalled RFs. (**A**) Immunoblots showing depletion of lamin-A/C in HEK293T and MCF7 cells lentivirally transduced with shRNAs directed against *LMNA* gene (shLmna) or luciferase gene (shLuc) as control. Similar levels of Lamin B1 expression were observed. (**B**) Single-molecule replication analysis performed in HEK293T cells generated in (A). Images show DNA fibers in cells labeled with IdU 20 min + CldU 20 min as detected by fluorescence confocal microscopy. Tract lengths are measured using ImageJ. Graph shows the tract length ratio CldU/IdU in untreated cells (NT), cells treated with hydroxyurea to stall RFs (4 mM HU for 3 hours), and cells treated with HU and the MRE11 nuclease inhibitor Mirin (50 μM). Average ± SEM of 3 biological repeats (3 independent lamin A/C depletions) is shown, with ~300 forks measured per experiment. Note how the ratio CldU/IdU<1 in shLmna cells treated with HU is rescued by Mirin. (**C**) Immunoblots show depletion levels of MRE11 nuclease in cells generated in (A) and transiently transfected with siRNA directed against MRE11 (siMRE11) or negative control (siCtrl). Histone 3 is the loading control. Graph shows tract length ratio CldU/IdU in cells not treated (NT) or treated with HU. Graph represents average ± SEM of 2 biological repeats (2 independent MRE11 depletions), with ~200 forks measured per experiment. (**D**) Tract length ratio CldU/IdU in MCF7 depleted of lamins and treated with HU ± Mirin as in (B). (**E**) Immortalized MEFs from *Lmna^+/+^* and *Lmna^−/−^*were treated with HU ± Mirin as in (B) and CldU/IdU ratio calculated. Graph represents average ± SEM of 2 different lines of MEFs. Immunoblots show lack of expression of lamin A/C, but not lamin B1, in MEFs from *Lmna^−/−^* mice compared to *Lmna^+/+^.* Vinculin and Histone 3 are loading controls. (**F**) Tract length ratio CldU/IdU in HEK293T generated in (A) and either not treated (NT) or exposed to camptothecin (100 nM CPT) or aphidicholin (200 nM APH). Graph shows 2 biological repeats, with ~200 fibers measured per experiment. (**G**) Cells generated in (A) were lentivirally transduced with either a mutant lamin A (LAΔ50 or progerin), wild type lamin A (LA), or an empty vector (EV) control. Immunoblots show levels of expression of progerin and lamin A. Graph shows ratio CldU/IdU in these cells either untreated (NT) or exposed to HU. Average ± SEM from 3 biological repeats is represented, with ~100 fibers measured in each experiment. All DNA fiber assays were performed with the same labeling scheme as in (A). In all figures, * denotes p value of statistical significance (*p<0.05; **p<0.01; ***p<0.001; ****p<0.0001), and error bars represent standard error of the mean (SEM).

Interestingly, the decreased CldU/IdU ratio upon RF stalling and the rescue of replication defects by Mirin were also observed in lamins-depleted tumor cells – MCF7 (**Fig 1D**) and U2OS (**Fig S3**)-, and in MEFs from *Lmna^−/−^* mice (**Fig 1E**). These data indicate that lamins are required for the stability of RFs in multiple cell types. Moreover, compounds such as camptothecin and aphidicolin, which cause RF stalling via different mechanisms, also elicit RFI in lamins-depleted cells (**Fig 1F**). Importantly, reconstitution of lamin A (LA) in lamins-depleted cells rescues replication defects upon HU treatment (**Fig 1G**). In contrast, expression of a truncated mutant form of lamin A (LAΔ50), also known as “progerin”, does not rescue replication defects. Rather, progerin itself imposes a challenge to DNA replication, causing RFI (reduced CldU/IdU ratio) in the absence of any drug. This confirms our published results showing fork stalling and nuclease-mediated degradation of newly replicated DNA upon expression of progerin in normal cells (49). In summary, our data demonstrate that lamins are involved in the protection of the replication intermediates that are formed at stalled RFs, with loss of lamins function mirroring RFI phenotypes of HR-deficient cells.

### Lamins loss elicits replication stress-induced genomic instability

Inability to maintain fork stability in HR-deficient cells results in the accumulation of DSBs and chromosomal aberration in response to RS (26,27,32,50). To determine if nucleolytic degradation of stalled forks causes genomic instability in lamins-deficient cells, we hindered RF progression by HU treatment and monitored the generation of DSBs by neutral comet assays (**Fig 2A**), and chromosomal aberrations in metaphase spreads prepared right after induction of replication stress by colcemid (**Fig 2B**). With this approach, increased DSBs and chromosomal aberrations over basal levels should derive from those RFs that stalled/collapsed during S-phase by HU treatment (27). As reported, lamins-deficient cells exhibit higher basal levels of DSBs and chromosomal aberrations than control cells (51). Prolonged exposure to high HU dosage induces fork degradation and accumulation of DNA damage (DSBs) and chromosomal breaks in control and lamins-deficient cells, in agreement with RS-induced DNA damage. Consistent with the notion that DNA damage can derive from nucleolytic degradation of unstable forks, inhibition of MRE11 with Mirin reduces DSBs and chromosomal aberrations in HU-treated lamins-deficient cells, with no effect in control cells (**Fig 2A,2B**). Interestingly, we did not observe complex chromosomal aberrations (i.e. triradials, dicentric chromosomes) typical of cells deficient in HR factors such as BRCA1/2 undergoing replication stress (26,27,32,50). This suggests that in lamins-deficient cells, genomic instability could derive from degradation of replication intermediates that differ from those arising in BRCA-deficient cells, despite the RFI phenotype mirroring BRCA deficiency. Overall, our results suggest that excessive remodeling of stalled forks is toxic in lamins-deficient cells and contributes to genomic instability. As evidence for lamins loss hindering the ability of cells to cope with replication challenges, we find that tumor cells (MDA-MB-231) depleted of lamins exhibit increased sensitivity to replication inhibitors, including HU and the poly(ADP-ribose) polymerase 1 (PARP1) inhibitor olaparib (**Fig 2C**). Increased sensitivity to olaparib is consistent with lamins-depleted cells exhibiting a replication phenotype that resembles HR-/BRCA-deficient cells.

**Figure 2.**
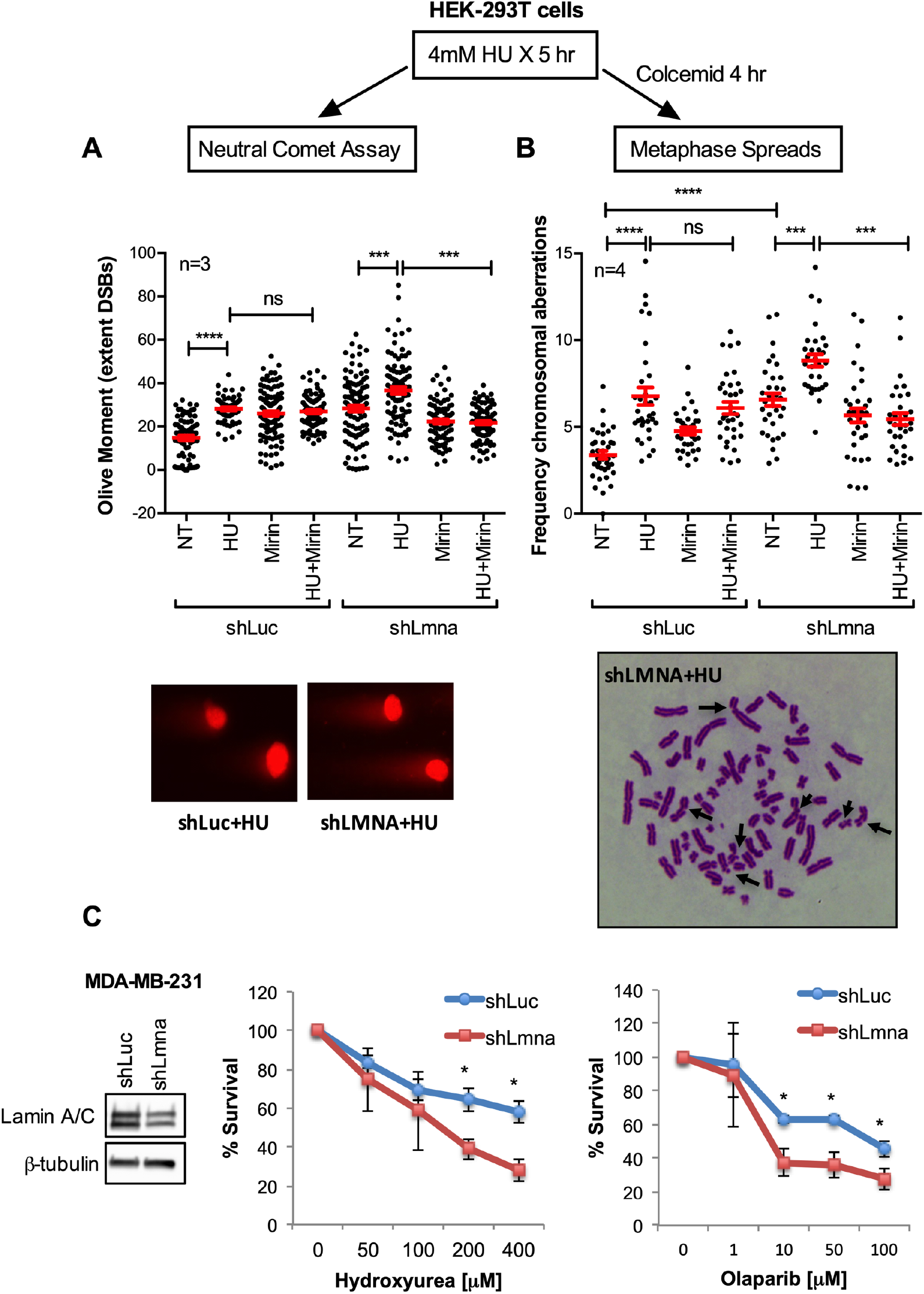
Replication stress causes genome instability in lamins-deficient cells. HEK293T cells depleted of lamins (shLmna) or control cells (shLuc) were either not treated (NT) or exposed to HU (4 mM), Mirin (50 μM), or the combination HU+Mirin for 5 hours. (**A**) After the treatments, cells were processed for Neutral Comet Assay to monitor the amount of DNA DSBs generated in response to RS. Graph shows average ± SEM of DNA DSBs in 3 biological repeats. (**B**) After the treatments, cells were incubated in 10 μM Colcemid for 4 hours and metaphase spreads prepared to monitor the frequency of chromosomal aberrations induced by RS (chromosome breaks, chromatid breaks, chromosome end-to-end fusions, and other complex aberrations). Graph shows average ± SEM of 4 biological repeats. Note how the increased genomic instability induced by HU is rescued by Mirin in lamins-deficient cells by both assays. (**C**) Graphs show survival curves of MDA-MB-231 cells depleted for lamin A and either left untreated or treated with increasing concentrations of HU or the PARP inhibitor Olaparib. Lamins-deficient cells exhibit increased sensitivity to both drugs in 3 biological repeats.

### Lamins are required for RAD51 protective function at stalled replication forks

Our results show that lamins are required to stabilize damaged RFs and to prevent genomic instability during RS. Stabilization and protection of stalled forks are essential processes during fork remodeling, a complex mechanism orchestrated by a multitude of factors that cooperate to reinstate DNA replication and to prevent that damaged forks translate into DSBs (31). BRCA2 plays an important fork protective role by mediating the loading of RAD51 on regressed arms (26,27). Interestingly, BRCA2 physically interacts with lamins during meiotic recombination in Drosophila oocytes (52). However, whether these two proteins also interact during DNA replication remains unknown. Lamins-depleted cells exhibit similar replication defects as BRCA2-deficient cells (**Fig 1**), suggesting that lamins and BRCA2 might be part of the same pathway. To test this hypothesis, we generated HEK-293T cells depleted of both, lamins and BRCA2 (**Fig 3A**) and performed DNA fiber assays labeling with thymidine analogs for twenty, thirty, and forty minutes to ensure that the length of green tracts was adequate for accurate measurements. BRCA2 depletion did not further exacerbate the RFI phenotype of lamins-depleted cells (**Fig 3B** and **Fig S4**), suggesting that lamins and BRCA2 act on the same RF protective pathway. Consistently, Mirin treatment rescued replication defects to the same extent in all three cell lines (**Fig 3C**). Importantly, lamins depletion does not affect BRCA2 expression, and thus lamins might hinder the RF protective function of BRCA2 or other factors in the pathway.

**Figure 3.**
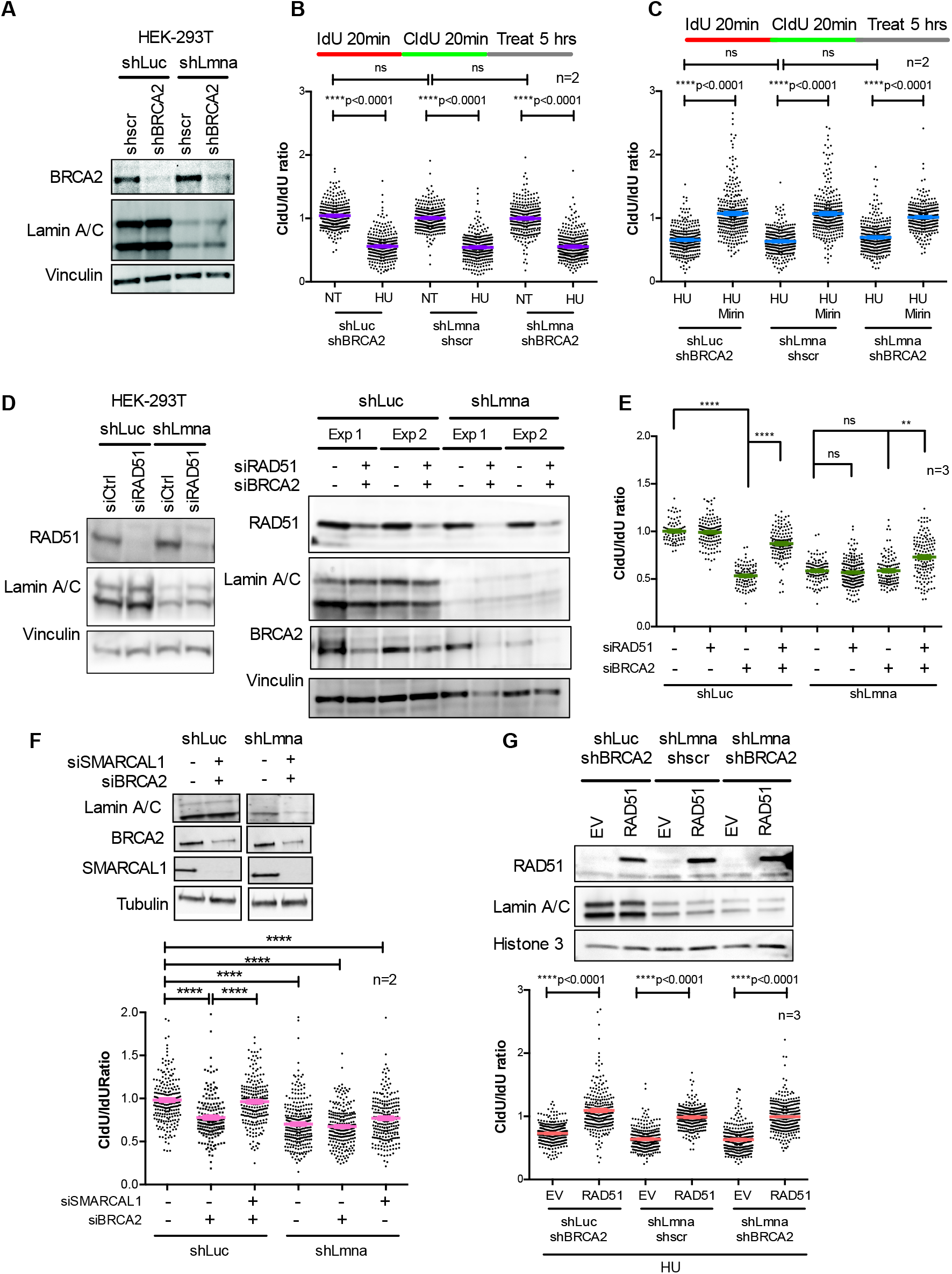
Replication defects in lamins-deficient cells have similarities with BRCA2-deficient cells. **(A)** HEK293T cells depleted of lamins (shLmna) or control cells (shLuc) were lentivirally transduced with short hairpin directed against BRCA2 or a scrambled control (shscr). Immunoblots show reduced levels of BRCA2 and lamin A/C proteins. Vinculin serves as loading control. (**B**) Singlemolecule replication analysis performed in HEK293T cells generated in (A). Cells were labeled with IdU 20 min + CldU 20 min and left untreated or exposed to 4 mM HU for 5 hours. Graph shows average ± SEM of tract length ratio CldU/IdU in cells generated in (A) and either left untreated (NT) or treated with HU. Cells depleted of BRCA2, lamin A/C, or both proteins share similar phenotypes under replication stress (CldU/IdU<1). Data from 3 independent experiments from 2 biological repeats, with ~200 fibers measured per experiment. (**C**) Cells generated in (A) and labelled as in (B) are exposed to HU alone or in combination with Mirin and used in single-molecule replication analysis. Graph shows tract length ratio CldU/IdU. Note how inhibition of MRE11 rescues fork degradation similarly in all the three different lines. Data from 3 independent experiments from 2 biological repeats, with ~200 fibers measured per experiment. (**D**) Immunoblots showing siRNA-mediated depletion of RAD51 (left panels) and BRCA2 and RAD51 (right panels) in lamins-proficient and -deficient cells. Vinculin is the loading control. Representative image of three biological repeats. (**E**) Graph shows tract length ratio CldU/IdU of cells generated in (D) and labelled with IdU 20 min + CldU 20 min prior to exposure to HU for 5 hours. Note how RAD51 depletion rescues RFI in BRCA2-deficient cells but not in lamin A-deficient cells. (**F**) Immunoblots showing lamins-proficient and – deficient cells transfected with siRNAs for depletion of BRCA2 and SMARCAL1. Graph shows tract length ratio CldU/IdU of lamins-proficient and -deficient cells depleted of SMARCAL1 and/or BRCA2 via siRNAs. Note how SMARCAL1 depletion rescues resection in BRCA2-deficient cells but not in lamins-deficient cells. (**G**) Cell lines generated in (A) were transduced with RAD51 construct or an empty vector (EV) control. Immunoblots show the overexpression of RAD51 in the three cell lines. Graph shows the ratio CldU/IdU in the six different cells. Note how overexpression of RAD51 rescues replication defects in all three lines. Data from three biological repeats and ~200 fibers measured in each experiment.

Next, we interrogated the involvement of RAD51, which plays a dual role at stalled RFs: promoting fork reversal independently of BRCA and protecting the regressed arms from nucleolytic degradation in a BRCA-dependent manner (11). Accordingly, depletion of RAD51, and thus inhibition of fork reversal alleviates nucleolytic degradation in BRCA2-deficient cells (28,33). Paradoxically, overexpression of RAD51 also inhibits nucleolitic degradation in BRCA2-deficient cells (24,26,27). This protective function of RAD51 is exerted through the formation of stable nucleoprotein filaments at regressed arms, an ability that is diminished in the absence of BRCA2 and restored by forced expression of RAD51 (34,36). To define how lamins loss impacts all these RAD51 protective roles, we first tested whether unprotected regressed arms are the entry point for MRE11 in lamins-deficient cells. In lamins-deficient and -proficient cells, we depleted RAD51 (**Fig 3D**) and monitored fork stability by DNA fiber assay (**Fig 3E**). As expected, depletion of RAD51, BRCA2, or both combined does not affect normal RF progression (**Fig S4D**). Consistent with previous studies (28,33), RAD51 depletion in BRCA2-deficient cells rescues fork resection (**Fig 3E**). Unexpectedly, depletion of RAD51 in lamins-deficient cells does not prevent fork resection. Similar results were obtained in two different cell lines depleted of SMARCAL1 (**Fig 3F** and **Fig S5A**), a fork remodeler involved in fork reversal and shown to contribute to RFI in BRCA2-deficient cells (32). The fact that in lamins-deficient cells, knockdown of RAD51 or SMARCAL1 does not rescue fork protection argues against the reversed fork being the point of entry of nucleases, implying that reversed forks are not always required for nucleases to resect nascent DNA. We did find however that overexpression of RAD51 (**Fig 3G**) prevents HU-induced fork resection in lamins-deficient cells, as well as in BRCA2-deficient cells and in cells depleted of both lamins and BRCA2 (**Fig 3G**). As control, we show that RAD51 overexpression does not impact normal RF progression in BRCA2-or lamins-depleted cells (**Fig S5**). Altogether, our results suggest that lamins deficiency hinders the protective function of BRCA2 and RAD51 at the RF. The defects however seem to be independent of fork reversal and more related to the inability to properly protect ssDNA generated during RF stalling.

### Lamins-deficient cells display delayed replication restart but normal fork reversal

The inability to rescue fork stability in lamins-deficient cells by depletion of factors involved in fork reversal (RAD51 or SMARCAL1) prompted us to investigate RF restart dynamics. Progressing forks are first labeled with IdU (red), then arrested with HU for 2 hours, and successively allowed to recover in complete medium supplemented with CldU (green), to label restarting forks (**Fig S1A**). This labeling scheme allows to distinguish stalled forks (red only) from restarting forks (red-green) and to identify newly fired origins (green only). Consistent with a normal cell cycle profile (**Fig S2**), lamins-depleted cells exhibit a similar percentage of progressing and spontaneously stalled forks as control cells (**Fig 4A**). As expected, in both cell lines there is an increased frequency of stalled forks in response to HU treatment and upon inhibition of MRE11 (HU+Mirin) (**Fig 4A**), as controlled nucleolytic resection of regressed arms is required to reinstate DNA replication (Mason et al., 2019; Delamarre et al., 2020). To survive RS, human cells fire dormant origins within active replicon clusters (X.Q. Ge et al., 2007). Interestingly, in lamins deficient cells the percentage of new origin fired in response to HU is significantly lower than in control cells (**Fig 4B**). This result is difficult to explain but given the complex nature of DNA replication initiation we cannot exclude a role of lamins in origin specification/licensing during G1 or suboptimal functioning of S-phase checkpoints. Unlike previous observation (16), we found that in lamins-deficient cells the proportion of forks restarting replication upon HU treatment is not statistically different from control cells. However, lamins loss results in delayed restart, with shorter CldU tracts compared to control cells (**Fig 4C**), similarly to what was previously reported in BRCA2-deficient cells (50). Controlled resection of ssDNA at regressed arms is necessary to restart replication while excessive nucleolytic activity is toxic. Interestingly, inhibition of MRE11 does not rescue timely restart of stalled forks in lamins deficient cells (**Fig 4C**). Our data argue that although lamins are dispensable to reinstate DNA replication, they play a role in ensuring timely restart of replication forks.

**Figure 4.**
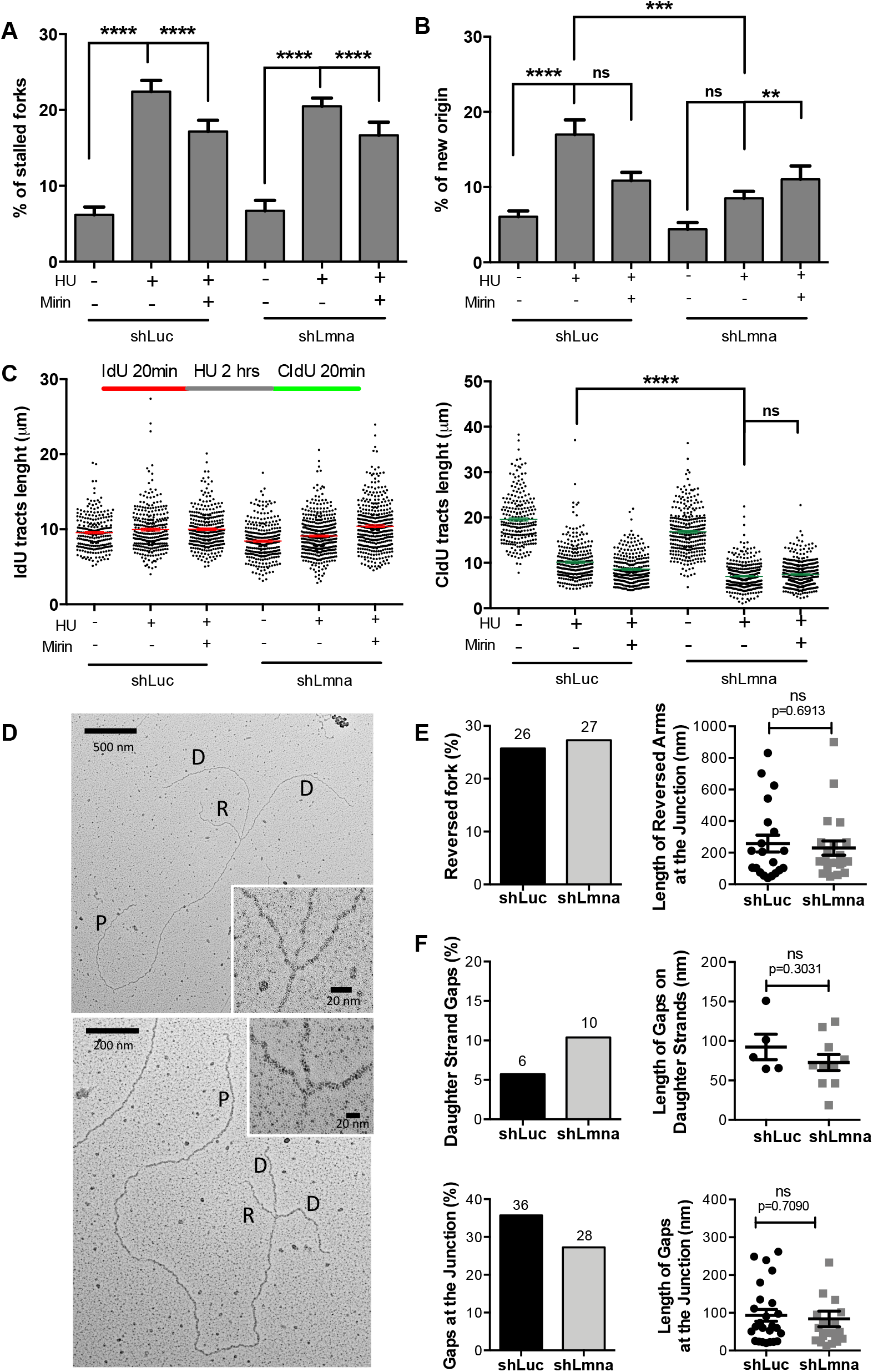
RFI in lamins-depleted cells is independent of fork reversal. (**A**) Immunoblots showing transient depletion of RAD51 (siRAD51 left panel) and BRCA2 (siBRCA2 right panel) via short interfering RNA in HEK293T cells proficient (shLuc) or deficient for lamins (shLmna). Two independent combined depletions are shown in the right panel. (**B**) Cells generated in (A) were processed for DNA fiber assays. Graph shows tract length ratio CldU/IdU of generated cells exposed to HU. Data from 3 biological repeats (3 independent depletions). For each experiment, we measured ~100 fibers. (**C**) Immunoblots showing the transient depletion of SMARCAL1 and BRCA2 via siRNAs in HEK293T cells proficient (shLuc) or deficient for lamins (shLmna). (**D**) Graph shows the result of DNA fibers (CldU/IdU ratio) performed in the different cells generated in (C). Note the RFI in BRCA2- and lamins-deficient cells and how depletion of SMARCAL1 only rescues the former. (**E**) Electron Microscopy performed in lamins-proficient and -deficient cells to visualize reversed forks and extent of ssDNA gaps at the stalled fork. The cells were treated with HU to induce RS and RFI in lamins-depleted cells and with Mirin to prevent degradation of reversed forks. RF structures are identified as two arms of equal length, and reversed forks as four-way junction structures. Two examples of reversed forks in lamins-depleted HEK293T cells are shown. (**F**) Frequency of fork reversal in lamins-proficient and -deficient cells treated with HU and Mirin (left graph) and length (nm) of the reversed arms at the junction. Note that there are no differences between control (sLuc) and lamins-deficient (shLmna) cells. (**G**) Frequency of ssDNA gaps in daughter strands and at the junction (left graphs) and length of the gaps (right graphs) in lamins-depleted and control cells.

Next, we investigated whether delayed fork restart was caused by an inefficient fork reversal process. To test this directly, we performed electron microscopy (EM), the only technique to directly visualize replication intermediates, including RFs that “reverse” their course following RS (28). We crosslinked cells with psolaren to preserve DNA structures and visualized by EM duplex DNA (10nm fiber) and ssDNA gaps (thickness 5-7nm). RF structures were identified as two arms of equal length, and reversed forks as four-way junction structures (**Fig 4D**). We compared lamins-proficient and – deficient cells subjected to HU treatment to stall the fork and to Mirin to prevent degradation of the newly synthesized DNA in the stalled fork. Interestingly, the frequency of fork reversal (**Fig 4E**) was indistinguishable between lamins-depleted and control cells. In addition, the frequency and length of ssDNA gaps in the daughter strands and at the junction were similar in lamins-depleted and control cells (**Fig 4F**). Thus, lamins loss does not have an impact on the generation of ssDNA at the stalled forks or the ability of forks to reverse as a mechanism of RF protection.

Altogether, our data suggest that lamins’ main role at stalled forks is to mediate the stability of RAD51 nucleoprotein filaments while having a minor role in fork reversal and restart. It is possible that in absence of lamins there is a limited recruitment of RAD51 to stalled forks such that fork reversal can still occur but protection of regressed arms is severely compromised thus leading to a slower restart. It is also possible that alternative factors such as RAD51 paralogs (RAD51B, RAC51C, RAD51D, XRCC2 and XRCC3) mediate RF reversal in lamins-deficient cells, as shown in other contexts (53,54).

### Lamins are essential for the recruitment of ssDNA binding proteins to stalled forks

Our data show that lamins-deficient cells are able to reverse stalled forks, yet they are severely impaired in protecting newly synthesized DNA from nucleolytic attack. Degradation of nascent DNA in lamins-deficient cells does not seem to come from regressed arms, given that inhibition of fork reversal does not rescue degradation. Thus, we placed our focus on the initial steps of RF protection during RS, which involve the recruitment of the heterotrimeric RPA complex to ssDNA generated upon RF stalling. RPA, considered the first responder to RS, interacts with a large number of replication and repair proteins, and precedes RAD51 loading on replication intermediates (**Fig 5A**). We hypothesized that the fork protective role of lamins could involve binding to the RF and facilitating the recruitment and function of these factors during RS. First, we monitored the association of RPA, RAD51 and BRCA1/2 with asynchronous and S-phase chromatin by subcellular fractionation. Cells were synchronized in S phase by overnight treatment with aphidicolin and released for two hours in normal media to allow the repair of any DNA damage and the reinitiation of replication (**Fig 5B**). Then, we induced RF stalling by HU treatment for two hours, followed by fractionation. Flow cytometry analysis confirmed efficient S-phase synchronization by this protocol and effective EdU incorporation during release, indicating efficient restart of replication prior to HU treatment (**Fig S6**). We did not find changes in global levels of factors involved in RF protection and remodeling upon depletion of lamins in asynchronous or S-phase synchronized HEK-293T cells, except for the levels of P-RPA^S33^, a marker of RS, which was markedly increased in control S-phase synchronized cells under HU treatment, but much less in lamins-deficient cells (**Fig 5B, WCL**). Consistent with RS, we did find a robust increase in RPA, RAD51, and especially P-RPA^S33^ in S-phase chromatin of control cells treated with HU. In contrast, lamins-depleted cells show markedly reduced levels of all these factors at chromatin upon RS (**Fig 5B, Chromatin S-phase**). These findings indicate that lamins facilitate the recruitment of key HR and fork remodeling factors to chromatin in response to RS. The same robust increase in P-RPA^S33^ levels was observed by immunofluorescence (IF) in control HEK-293T cells treated with HU (**Fig 5C**). In contrast, lamins-depleted cells show markedly reduced levels of P-RPA^S33^, supporting the idea that lamins loss hinders the proper recruitment of RPA to chromatin during RS, thus preventing phosphorylation. Accordingly, forced expression of lamin A in lamins-depleted cells rescued the levels of P-RPA^S33^, indicating that this is a direct consequence of lamins loss (**Fig 5C**). As control, we calculated the percentage of cells in S-phase by FACS (**Fig 5C, table**), which is similar in all conditions, thus implying that the differences in RFU are not due to changes in cell cycle. Importantly, the results are not cell-specific, as we confirmed in MCF7 cells (**Fig 5D**). Note how the robust increase in P-RPA^S33^ levels in control MCF7 cells treated with HU is prevented by lamins depletion. Altogether, these studies support the idea that lamins loss hinders the proper recruitment of RPA to chromatin during RS, thus preventing phosphorylation and proper RS signaling.

**Figure 5.**
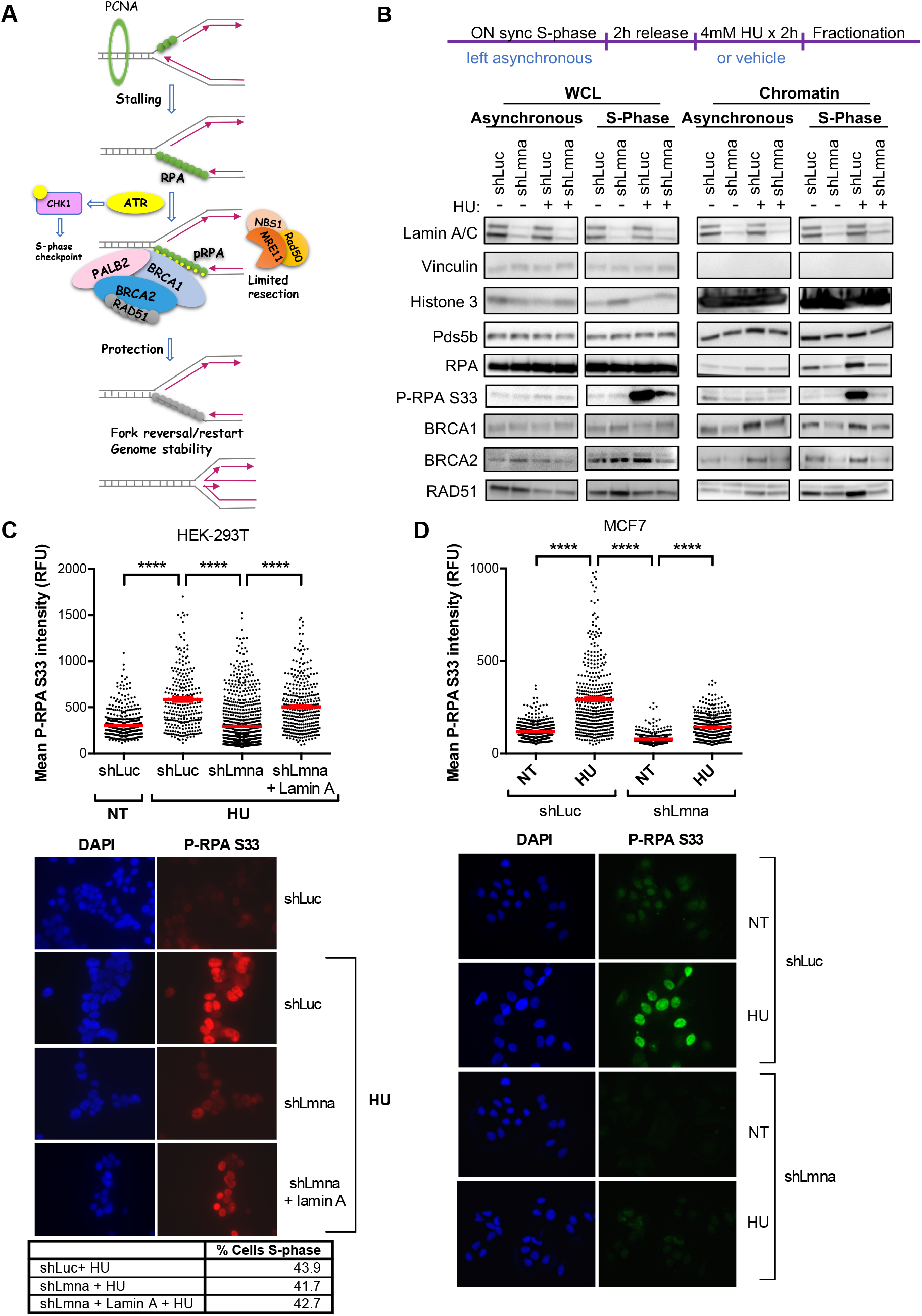
Lamins deficiency hinders chromatin recruitment and phosphorylation of RPA. (**A**) Schematic representation of the events occurring upon stalling of RFs. The uncoupling of the replicative polymerase from the helicase generates ssDNA that is coated by the ssDNA-binding trimeric complex RPA. This activates the ATR/CHK1 intra S-phase checkpoint and ATR-mediated phosphorylation of DNA-bond RPA, which becomes a docking site for a plethora of fork-protective/remodeling factors (BRCA1, PALB2, BRCA2 and RAD51). These factors stabilize the fork and mediate restart of the stalled fork. (**B**) HEK293T cells proficient (shLuc) and deficient for lamins (shLmna) were either left asynchronous or synchronized in S-phase via overnight treatment with 1 μM aphidicholin (APH). Cells were then allowed to recover for 2 hours in complete medium, followed by exposure to HU (4 mM) to elicit RS or vehicle as control for 2 hours. Subsequently, cells were either lysed directly to prepare whole cell lysates (WCL) or subjected to subcellular fractionation to isolate the chromatin fraction. Immunoblots show global protein levels from WCL or chromatin-bound proteins from asynchronous and S-phase synchronized shLuc and shLmna cells. Note how loss of lamins results in a reduction of proteins associated to S-phase chromatin under conditions of RS. Vinculin is the loading control for WCL, whereas Pds5b and H3 are loading controls for the chromatin fraction. Representative immunoblots of 3 biological repeats. (**C**) Immunofluorescence (IF) performed in HEK293T lamins-proficient (shLuc) and -deficient (shLmna) cells, as well as in cells in which lamin A expression was reconstituted (shLmna+lamin A), and treated with HU to induce RS. Monitored levels of P-RPA^S33^, a marker of RS that indicates binding of RPA to chromatin and phosphorylation by AKT. Representative fields of cells are shown in the images and quantification of fluorescence intensity (relative fluorescence units) is shown in the graph (ImageJ program). DAPI staining was used to demarcate nuclei. Results are from 2 independent experiments and >200 cells quantified per condition each time. Table shows percentage of cells in S-phase by FACS. (**D**) Lamins-proficient and -deficient MCF7 cells were processed for IF as in (C) to monitor the levels of P-RPA^S33^ in response to HU treatment. Note the marked decrease in fluorescence intensity in lamins-deficient cells compared to control. Results are from 2 independent experiments.

Because subcellular fractionation and immunofluorescence do not allow to distinguish global chromatin from replication forks, we performed iPOND to monitor directly the binding of proteins to nascent DNA (**Fig 6A**). HEK-293T cells were pulse-labeled with EdU for fifteen minutes and either left untreated (NT) as control, subjected to a sixty minutes Thymidine chase, or treated for two hours with HU, which is sufficient to stall RFs without causing DNA DSBs (**Fig S7**). “Input” samples show similar levels of proteins in these cells under the different conditions, except for P-RPA and γH2AX, which increase upon HU treatment due to RS and DNA damage (**Fig 6A**). This is in line with results in **Fig 5B** showing that the global levels of proteins are not affected by lamins loss. “Capture” samples show binding of lamin-A to nascent DNA (NT) in control (shLuc) cells consistent with previous observation by nascent chromatin capture proteomics (55). Importantly, we find that lamins association to nascent DNA increases upon fork stalling (HU) supporting the notion that these nuclear proteins assist damaged RFs. As expected, HU treatment results in increased binding of RPA, P-RPA and RAD51 in control cells, indicating a proper response to fork damage. However, lamins-depleted cells exhibit a robust shortage of binding of RPA, P-RPA and RAD51 to nascent DNA (**Fig 6A**). Histone H3 (loading control) and γH2AX (DNA damage marker) are associated with nascent DNA to the same extent in lamins-proficient and -deficient cells, excluding the possibility that lack of protective factors at stalled forks in lamins-deficient cells is due to differences in loading or to ineffective HU treatment.

**Figure 6.**
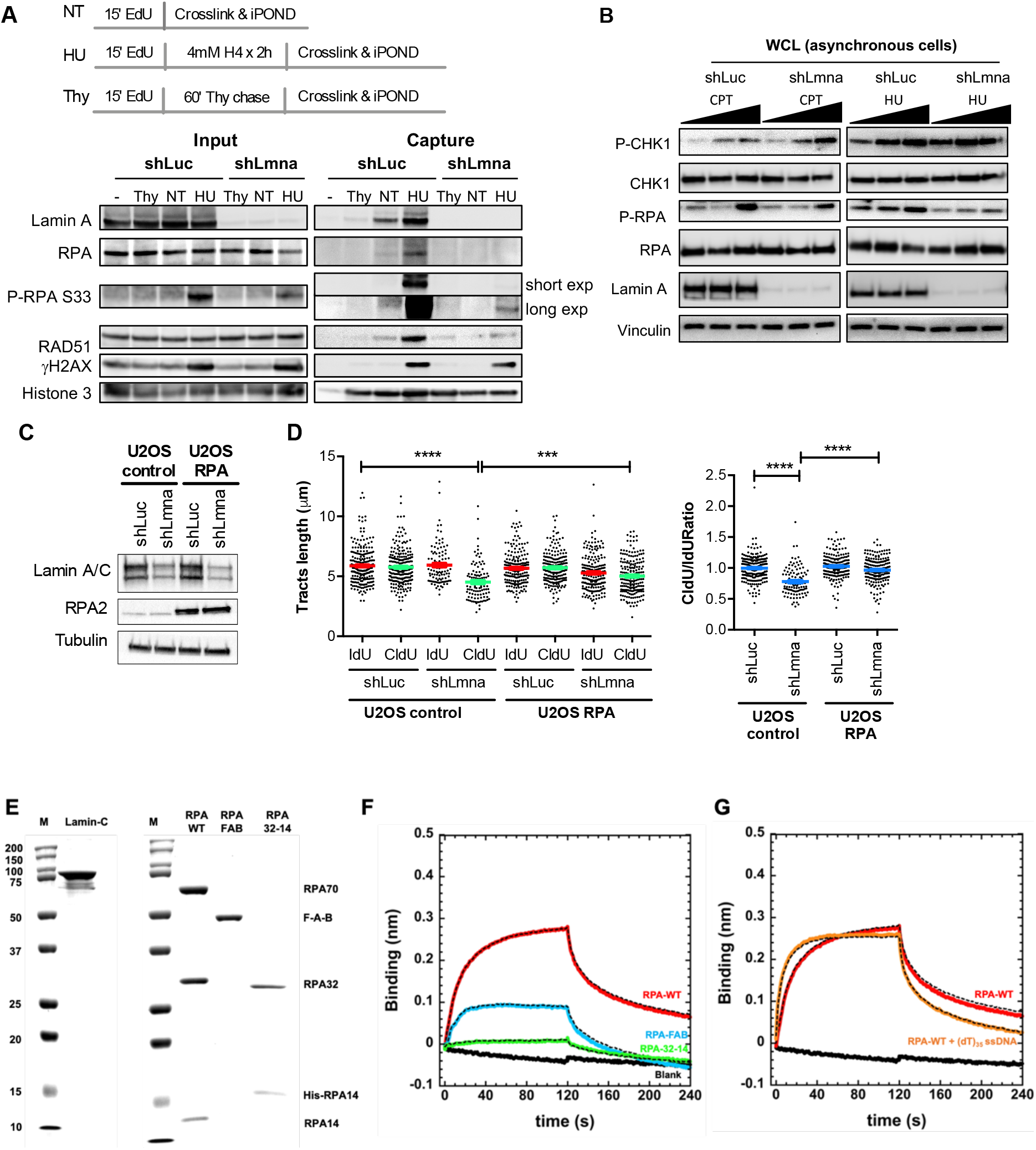
Lamins interact with RFs and RPA and are required to recruit RPA and RAD51 to stalled forks. (**A**) iPOND of shLmna and shLuc cells under basal conditions (NT) and upon replication stress (HU). HEK293T shLuc and shLmna cells were EdU-labeled and treated with 4 mM HU for 2 hours. Thymidine chase experiment is performed to distinguish proteins associated with post-replicative chromatin from those at active RFs. Input (2%) shows global levels of proteins in the starting material. The immunoblot of captured proteins shows that lamins directly associate with RFs and this interaction increases under replication stress (HU). Note the marked reduction of RF protective factors (RPA, P-RPA, RAD51) at active forks in lamins-deficient cells. (**B**) Immunoblots showing levels of CHK1 and RPA, and activation of both proteins by phosphorylation (P-CHK1 and P-RPA) in response to RS induced by HU or camptothecin (CPT) treatment for 2 hours. Increasing concentrations of CPT (25 nM, 100 nM, 1 μM) or HU (0.5 mM, 1 mM, 2 mM) were used. Note how in absence of lamins global levels of P-RPA are reduced. Vinculin is used as loading control. Representative immunoblot of 3 biological repeats. (**C**) U2OS cells control and super-RPA U2OS cells were transduced with shRNAs for stable depletion of lamins (shLmna), and with shLuc as control. Immunoblots show the reduced levels of lamin A/C and the overexpression of a subunit of the RPA complex (RPA2). (**D**) The graphs show the results of DNA fiber assays performed in the four different cell lines. Note the shortening of the green tracts (left graph) and the ratio CldU/IdU<1 (right graph) in lamins-deficient U2OS control cells, which is not observed in super-RPA U2OS cells. (**E**) SDS-PAGE analysis of recombinantly purified human Lamin-C, RPA, RPA-FAB and RPA32-14. (**F**) Biolayer interferometry (BLI) analysis of interactions between RPA or RPA sub-domains and Lamin-C show binding between the two proteins with an estimated K_d_ of ~35 μM. The FAB fragment of RPA shows binding to Lamin-C and very weak binding to RPA32-14 is observed. (**G**) Lamin-C binds with higher affinity when RPA is pre-bound to a (dT)_35_ ssDNA oligonucleotide (K_d_ ~1.5 μM).

These findings strongly suggest that lamins bind to nascent DNA and that facilitate the recruitment of ssDNA binding proteins during replication stress, thus explaining the excessive resection phenotype observed in lamins-deficient cells. While the protective role of RAD51 has been extensively studied, only recently it has been shown that P-RPA can directly inhibit DNA resection by limiting the processivity and speed of nucleases (56). We exclude the possibility that low P-RPA observed in lamins-deficient cells is due to a compromised ATR activity, as we found a dosedependent activation of Chk1, an ATR effector kinase activated in response to RS (57), to increasing concentrations of either camptothecin or HU (**Fig 6B**). Our results argue that failure to protect stalled forks is due to faulty recruitment of both RAD51 and RPA to nascent DNA. Being RPA the first responder to RS (48), we propose that limited recruitment of RPA to ssDNA plays a major role in the RFI observed in lamins-depleted cells. To test this model directly, we monitored RF stability by DNA fiber assay in a line of U2OS cells overexpressing the heterotrimeric RPA complex (super-RPA cells) (39) and transduced with shRNAs targeting lamins or control hairpins (**Fig 6C**). We show that in U2OS control cells under replication stress (HU treatment), depletion of lamins results in RFI, as shown by the shortening of the green tract (**Fig 6D, left graph**) and by a ratio CldU/IdU<1 (**Fig 6D, right graph**). In contrast, RFI is not observed in lamins-depleted cells that overexpress the RPA complex (**Fig 6D**). The prevention of RFI upon forced expression of the RPA complex in lamins-deficient cells indicates that sub-optimal recruitment of RPA at stalled forks might be the main cause of fork protection failure in lamins-deficient cells. Since RPA binding precedes RAD51 on nascent DNA, reduced amount of this “docking” factor could impact on RAD51 loading and therefore on the ability of cells to remodel stalled forks. Limited recruitment of these factors could also help explain the slow replication restart observed in lamins-deficient cells despite their preserved ability to perform fork reversal. Overall, our study indicates a key role for lamins in the protection of ssDNA from degradation during RS by mediating the recruitment of ssDNA binding proteins RPA and RAD51 to stalled RFs.

### Lamins physically interact with RPA

Since lamins influence the recruitment of RPA to ssDNA, we next tested whether the proteins interact. Lamin-C was recombinantly purified (**Fig 6E**) and binding to RPA was tested using biolayer interferometry (BLI). Lamin-C was tethered onto the optical probe and binding to RPA was captured (K_d_ = 35 μM; **Fig 6F**). When DNA is pre-bound to RPA, the affinity of this complex increases (K_d_ = 1.5 μM; **Fig 6G**). RPA binds to RPA interacting proteins through two separate protein-interaction domains or compositely through interactions with multiple regions in RPA. The F-domain in the large RPA70 subunit and the winged-helix (wh) domain in the RPA32 subunit are well-established protein interaction regions. To test whether Lamin-C interaction was coordinated through these domains, we tested Lamin-C binding to two fragments of RPA. The FAB fragment has the DNA binding regions (DBDs) A and B, and the F-domain. The other fragment of RPA had the RPA32 and RPA14 subcomplex (RPA32-14). In BLI experiments we qualitatively see physical interactions between Lamin-C and the FAB fragment but poor binding to RPA32-14 (**Fig 6F**). Thus, the F-domain likely is the major contributor to RPA interactions with Lamin-C.

## DISCUSSION

This study demonstrates a role of A-type lamins in DNA replication in mammalian cells. We propose a model whereby lamins are not essential for RF progression under unperturbed conditions but become critical under RS to protect stalled forks from the action of nucleases. Loss of lamins hinders the recruitment of RF protective and remodeling factors, especially ssDNA binding proteins RPA and RAD51, freeing the way for nucleolytic attack by MRE11. This function of lamins prevents genomic instability arising from challenges encountered during DNA replication.

An important finding is that lamins-depleted cells exhibit markedly reduced levels of P-RPA in response to RS. RPA phosphorylation, mediated primarily by ATR (58), is a critical early step in the loading cascade of protecting factors at stalled RFs (59). ATR accumulates at the nuclear envelope during S-phase (60). Thus, it is possible that lamins depletion hinders ATR localization at the nuclear periphery and potentially to other ssDNA-RPA entities located away from the nuclear envelope. There is also evidence that RPA directly interacts with MRE11 during unperturbed replication, and that P-RPA displaces MRE11 from this interaction (61–63). Deficiencies in P-RPA in lamins-depleted cells might result in MRE11 being retained at stalled RFs, where it could exert unrestricted nucleolytic activity. In addition, P-RPA is important for the recruitment of BRCA1/2 and PALB2 proteins to stalled forks (27,59), which in turn are required for the loading of RAD51 that promotes fork reversal and restart (29,35). Our findings that lamins-depleted cells exhibit deficiencies in RAD51 recruitment to nascent DNA are consistent with lamins loss hindering the whole cascade of events that protect stalled RFs. Interestingly, direct physical interaction between A-type lamins and BRCA2 has been previously reported (52), which regulates the association of chromatin flanking persistent DSBs along the nuclear envelope. Thus, it is also possible that A-type lamins directly mediate the positioning of BRCA2 at stalled forks. Our data showing that depletion of BRCA2 in lamin-deficient cells does not exacerbate replication defects observed upon dNTP depletion support the idea that lamins and BRCA2 cooperate to protect stalled forks. Depletion of either of these proteins would lead to deficiencies in RAD51 loading to stalled RFs.

The mechanisms whereby RAD51 stabilize the RF in response to RS has received much attention lately. Deficiency in the loading of RAD51 upon lamins loss is especially significant, as RAD51 is essential for the protection of the stalled forks as well as their remodeling via fork reversal, which facilitates proper restart of the RF (29,33). A direct interaction of lamin A with RAD51 has been reported (64), and thus a direct recruitment of RAD51 by lamins is possible. Interestingly, we find some differences in the effects of BRCA2-deficiency and lamins loss in DNA replication. The phenotype of BRCA2 deficient cells is rescued by both RAD51 overexpression and RAD51 depletion, as reported (26,28). A model to explain this paradoxical effect of RAD51 proposes that overexpression of RAD51 allows the protection of stalled and reversed forks from nucleolytic degradation. The rescue by RAD51 depletion is due to inhibition of fork reversal, the main point of entry of nucleases in BRCA2-deficient cells. In lamins-depleted cells, replication defects are only rescued by RAD51 overexpression, and not by RAD51 depletion. This suggests that fork reversal in lamins-deficient cells might be mediated by proteins other than RAD51 and that reversed forks in these cells do not constitute the main point of entry of nucleases.

Our study also shows that although RS does not compromise the viability of lamins-deficient cells, it results in accumulation of DSBs and chromosomal aberrations. This genomic instability might increase the susceptibility to acquire mutations in the genome that promote malignancy. For instance, decreased expression of A-type lamins has been observed in more than 30 cases of high-grade premalignant cervical epithelium lesions (65). Consistently, depletion of lamin-A/C in primary breast epithelial cells leads to cancer-like altered morphology and aneuploidy (66). Importantly, different subtypes of advanced breast cancers, including triple negative breast cancers (TNBC), feature low expression of A-type lamins, while high levels of these proteins are observed in normal mammary tissues (67,68). Perhaps the more complete study is one performed on a large well-characterized series of early stage operable breast cancers (938) with long-term follow-up (146 months on average), which showed that reduced or loss of expression of lamin A/C is associated with high histological grade, larger tumor size, poor Nottingham Prognostic Index (NPI, used to determine the prognosis following breast cancer surgery via pathological parameters such as size of the lesion, number of involved lymph nodes, and grade of the tumor), more distant metastasis, and shorter breast cancer-specific survival (69). Further studies are needed to determine the extent to which RFI upon lamins loss contributes to malignancy and cancer progression.

We propose that the role exerted by lamins in preventing the degradation of stalled forks and in the resultant genome instability has important implications for therapy. Depletion of A-type lamins in the TNBC cell line MDA-MB-231, harboring wild type BRCA1/2 proteins, increases sensitivity to HU and PARP inhibitors, which exploit defects in DNA replication and DSB repair. The increased sensitivity to PARP inhibitors is particularly interesting, as these compounds are at the forefront for the treatment of breast cancers that exhibit BRCA1/2 mutations. These results suggest that expression levels of A-type lamins might serve as a prognostic marker, similar to BRCA-deficiency, for breast cancer cells. Additional studies are also needed to determine whether the phenotypes observed in lamins-depleted cells are recapitulated in cells deficient in well-known lamin-binding proteins such as LAP2 isoforms (70) and Retinoblastoma (Rb) family (71), and whether other disease-associated *LMNA* gene mutations cause RS.

Collectively, these results advance our understanding of the role exerted by A-type lamins during DNA replication. In particular, we provide strong evidence for loss of lamins shifting the balance between protection and resection at stalled forks, resulting in genomic instability. These findings and the role of progerin eliciting RS suggest that special attention should be paid to the contribution of RS to genomic instability in different laminopathies. In a broader view, one could envision that while an intact nuclear lamina protects the genome of normal cells from the onset of mutations that trigger malignancy, disruption of this nuclear skeleton in cancer cells could increase the vulnerability of cancer cells to therapeutic compounds.

## ACKNOWLEDGEMENT

We acknowledge Denisse Carvajal and Annabel Quinet de Andrade for their help setting up the DNA fibers technology in our laboratory. S.Gr. and N.C.B. performed most of the experiments and S.Gr. wrote the first draft of the manuscript. B.T.C. and S.K. characterized the lamin C-RPA interaction in vitro by BLI, under the supervision of E.A., and E.S. and U.M. participated in some experiments. J.J. performed EM experiments. A.V. and E.A. provided insightful discussions throughout the study. We are in debt to Luis Toledo for sharing with us the super-RPA constructs. S.G. supervised the research and prepared the manuscript.

## FUNDING

Research in the S.G. laboratory is supported by NIA grant RO1AG058714. The work in the A.V. laboratory is supported by NIH grant R01GM108648 and by DOD BRCP Breakthrough Award BC151728. Work in the E.A. laboratory is supported by grants from the NIH GM130746 and GM133967. S.Gr. was recipient of a Pre-doctoral Fellowship from SCC. N.C.B. is recipient of a Postdoctoral Fellowship from American Heart Association.

## CONFLICT OF INTEREST

Authors declare no financial interests in relation to the work described.

## Notes

### Competing Interest Statement

The authors have declared no competing interest.

