## Supplemental figures for "A-type lamins are critical for the recruitment of RPA and RAD51 to stalled replication forks to maintain fork stability"

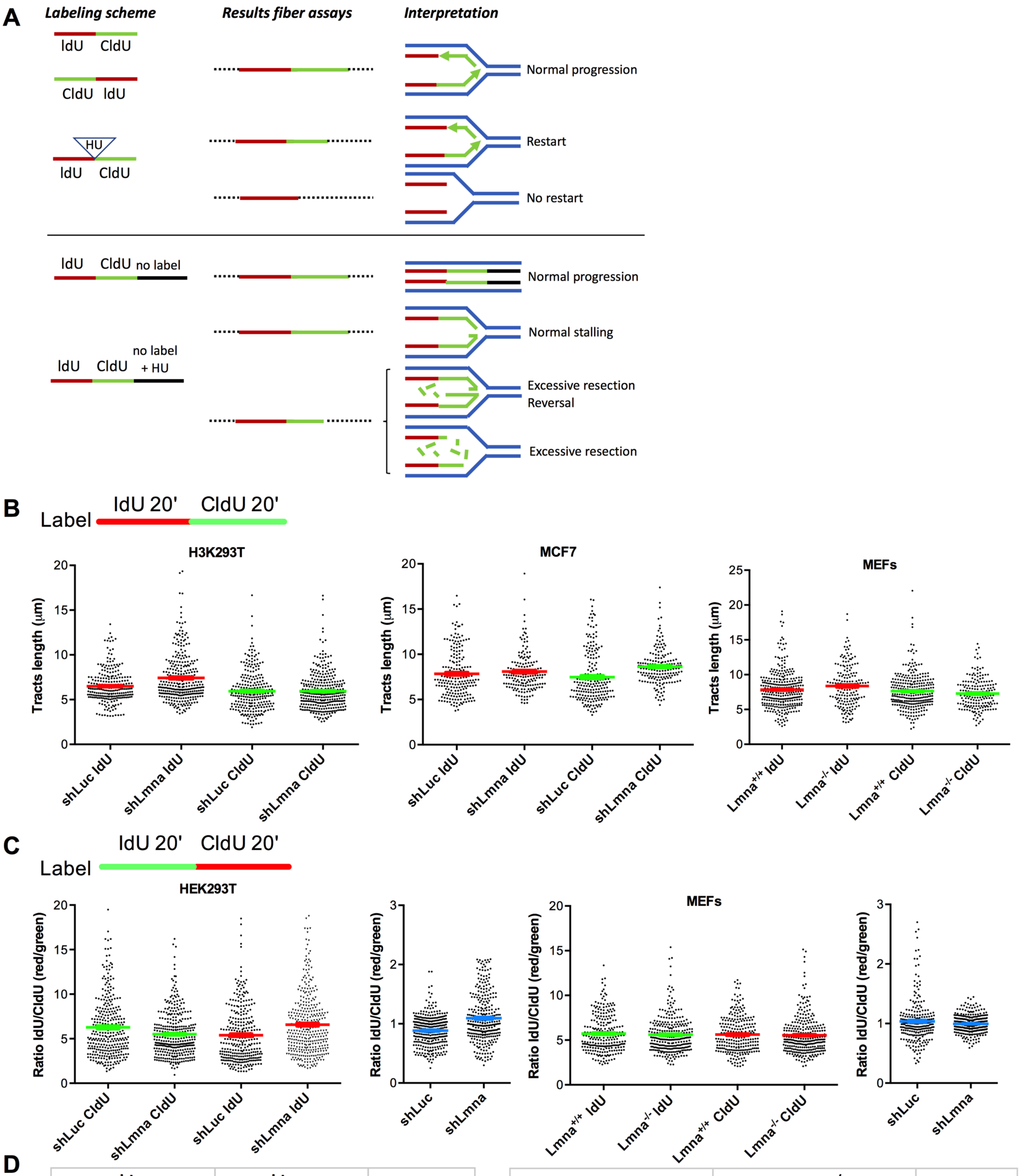


D

| shLuc | | shLmna | | |
|---|---|---|---|---|
| Green | Red | Green | Red | |
| 6.3 | 5.4 | 5.5 | 6.6 | AVERAGE µm |
| 3.5 | 3.2 | 2.5 | 3.5 | STDV µm |
| 0.2 | 0.2 | 0.1 | 0.2 | SEM µm |
| 16.3 | 14.0 | 14.2 | 17.0 | AVERAGE Kb |
| 9.0 | 8.2 | 6.4 | 9.2 | STDV Kb |
| 0.5 | 0.5 | 0.3 | 0.5 | SEM Kb |

| $Lmna^{+/+}$ | | $Lmna^{-/-}$ | | |
|---|---|---|---|---|
| Green | Red | Green | Red | |
| 5.7 | 5.6 | 5.6 | 5.5 | AVERAGE µm |
| 2.1 | 2.0 | 2.0 | 2.1 | STDV µm |
| 0.1 | 0.1 | 0.1 | 0.1 | SEM µm |
| 14.3 | 14.1 | 14.0 | 13.9 | AVERAGE Kb |
| 5.3 | 5.0 | 4.9 | 5.2 | STDV Kb |
| 0.3 | 0.3 | 0.3 | 0.3 | SEM Kb |

**Figure S1, related to Figure 1**. **DNA fiber assays in lamins-proficient and -deficient cells.** (**A**) Schematic representation of labeling protocols used to monitor different replication events by DNA fiber assays. Labeling with thymidine analogs is used in combination with drugs that slow-down or stall the RF in order to monitor rate of replication, RF stalling, resection, and fork restart. Different replication patterns and their interpretation are shown. (**B**) Lamins-proficient and -deficient cells (HEK293T, MCF7, and MEFs) were labeled 20 min with IdU, followed by 20 min with CldU, and DNA fiber assays performed in the absence of any drugs. The lengths of red (IdU) and green (CldU) tracts were measured by Image J and expressed as micrometer values. Graphs show average $\pm$ SEM of tracts length in biological repeats (3 for HEK293T cells, 2 for MCF7 cells, and 2 for MEFs). Note how red and green tracts show similar lengths in lamins-proficient and -deficient cells, suggesting that lamins loss does not alter normal replication fork progression. (**C**) DNA fiber assays were performed in HEK293T cells and in MEFs with the labeling inverted: 20 min CldU (green), followed by 20 min IdU (red). The IdU/CldU ratios are also shown. (**D**) Table shows the conversion of micrometers to kilobases in DNA fibers in (C) using a conversion factor: 1 $\mu$m=2.59 kb. Note how the rate of replication is similar in lamins-proficient and -deficient cells.

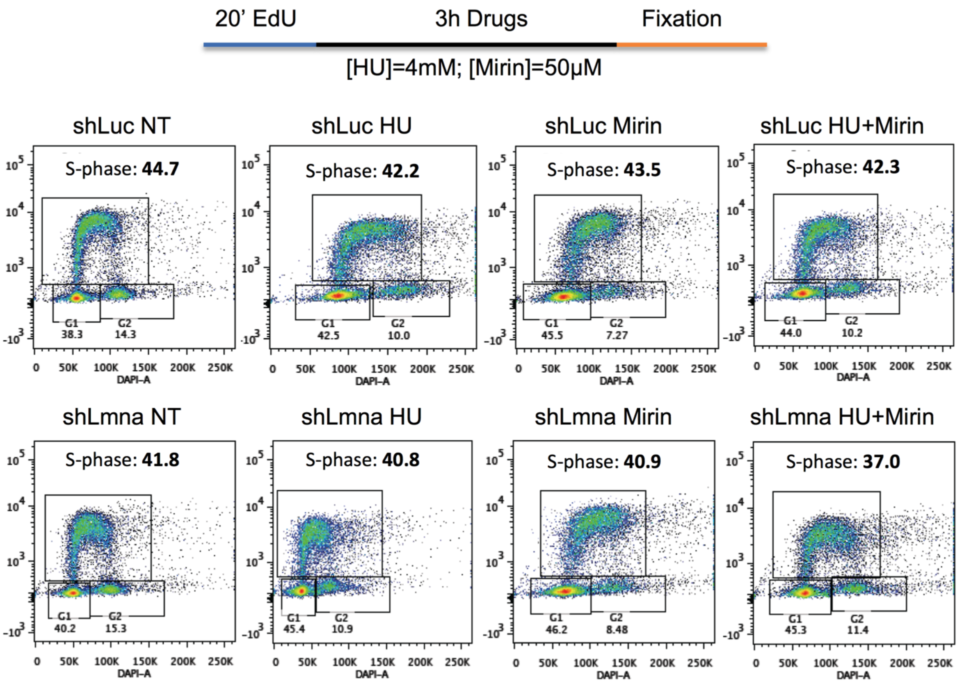


**Figure S2, related to Figure 1**. **Cell cycle profile of lamins-proficient and -deficient HEK293T cells subjected to different treatments.** HEK293T cells depleted of lamins (shLmna) and control (shLuc) were labeled with EdU for 20 minutes, followed by 3-hour treatments with HU, Mirin, or HU+Mirin. Cells were then fixed, permeabilized, and stained with DAPI. Cells were analyzed for fluorescent DNA content and EdU content using the BD Biosciences LSR II flow cytometer, and the cell cycle profiles were created by the program FlowJo®. Note how the percentage of cells in the S-phase of the cell cycle is similar in the different conditions, indicating that lamins loss does not affect progression through S-phase. Note also that while HU treatment does not significantly affect the percentage of cells in S-phase (shLuc 42.2% vs shLmna 40.8%), the distribution of shLmna cells is skewed to the left. We do not interpret this as a reflection of HU-induced accumulation of shLmna cells in early S-phase, but rather the reflection of extensive degradation of EdU-labeled nascent DNA at stalled forks. In line with this interpretation, EdU-labeled shLmna cells treated with HU+Mirin show a normal S-phase distribution. These results are consistent with the phenotype observed by fiber assay.

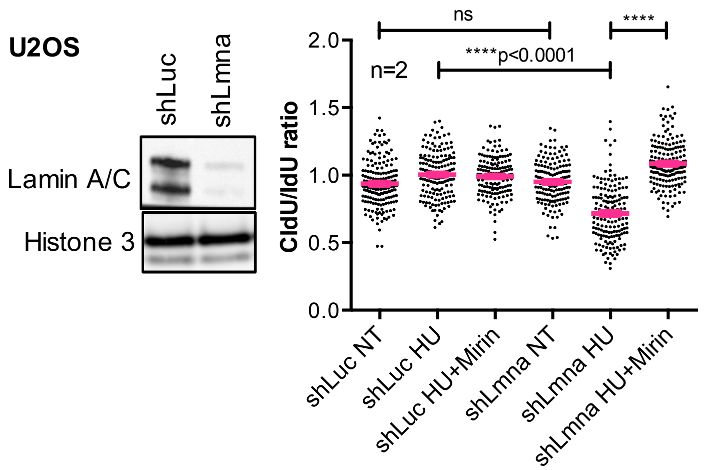


**Figure S3, related to Figure 1. Replication fork instability upon loss of lamins.** Immunoblots show depletion of lamin A/C in U2OS cells lentivirally transduced with shLmna or shLuc as control. Histone 3 is the loading control. Graph shows the results of DNA fiber assays performed in U2OS cells labeled with IdU 20 min + CldU 20 min. The tract length ratio CldU/IdU in untreated cells (NT), cells treated with hydroxyurea to stall RFs (4mM HU for 3 hours), and cells treated with HU and the MRE11 nuclease inhibitor Mirin (50mM). Note how the ratio CldU/IdU<1 in shLmna cells treated with HU is rescued by inhibition of MRE11. Graph shows average ± SEM of 2 biological repeats (2 independent lamin A/C depletions). In each experiment ~200 forks were measured. (**B**) DNA fiber assays performed in *LAP2*$\alpha^{+/+}$ and *LAP2*$\alpha^{-/-}$ MEFs (obtained from Roland Foisner) with the labeling scheme: IdU 20 min + CldU 20 min.

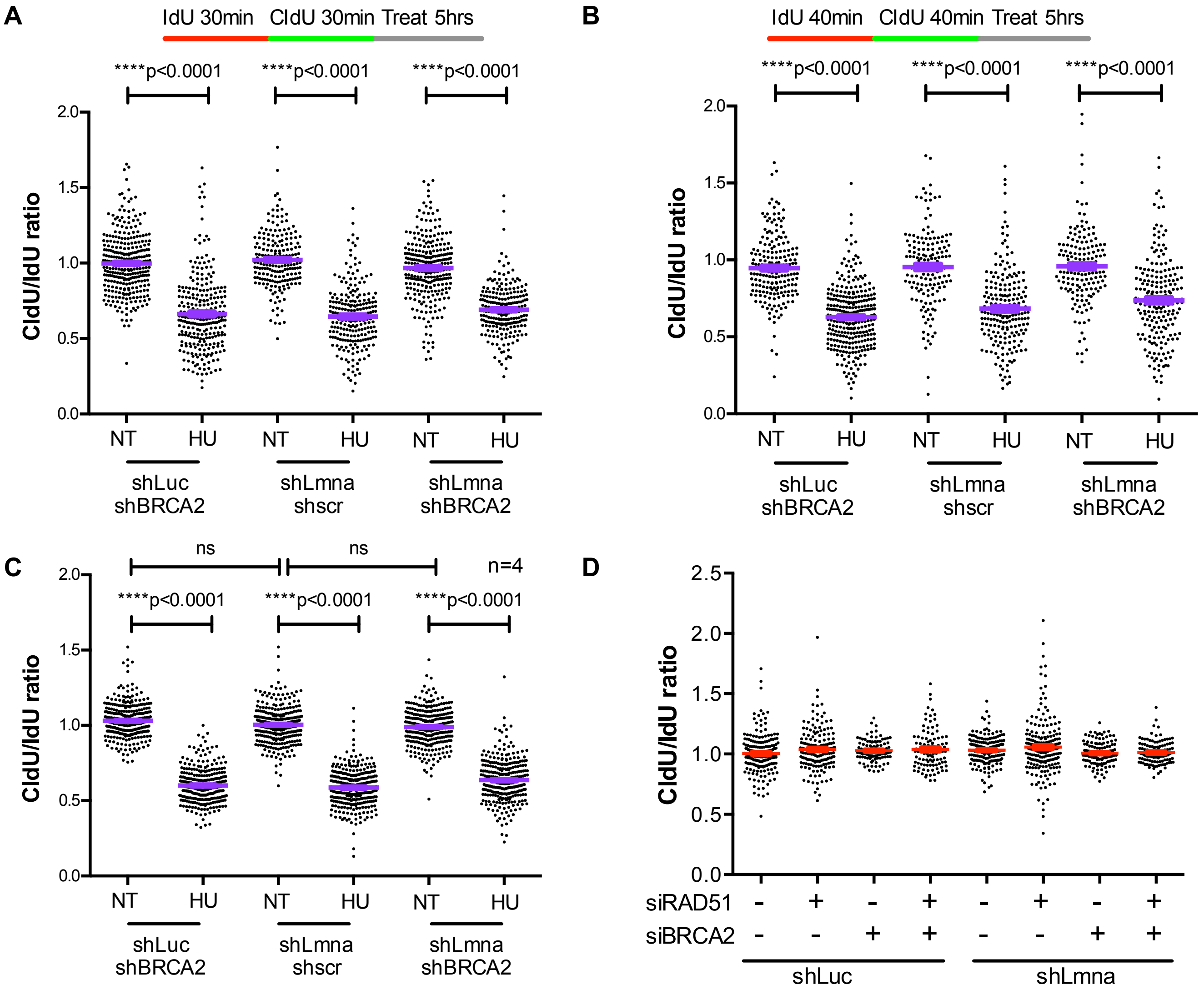


**Figure S4, related to Figure 3. Similar DNA replication defects in cells depleted of lamins, BRCA2, or both combined**. DNA fiber assays performed in the three cell lines with two different labeling schemes: (**A**) 30 min IdU + 30 min CldU, and (**B**) 40 min IdU + 40 min CldU. These labeling schemes were used to increase tract lengths, and thus rule out that the lack of differences among HU-treated cell lines is due to green tracts being too short for accurate measurements. Note that the same results are obtained with the different labeling schemes. (**C**) Graph shows the average ratio CldU/IdU of all the biological repeats (with the different labeling schemes) performed in cells depleted of lamins, BRCA2, or both combined. Note how BRCA2 depletion does not further exacerbate the fork degradation phenotype of lamins-deficient cells. (D) Graph shows tract length ratio CldU/IdU in HEK293T cells depleted of lamins, BRCA2, or both combined, and transfected with an siRNA targeting RAD51 or an siRNA control. Cells were labeled with IdU 20 min + CldU 20 min and not exposed to any drug treatment. Note how the ratio CldU/IdU is ~1 in all the cell lines and thus not affected by RAD51 depletion. Data from three biological repeats and ~100 fibers measured in each experiment.

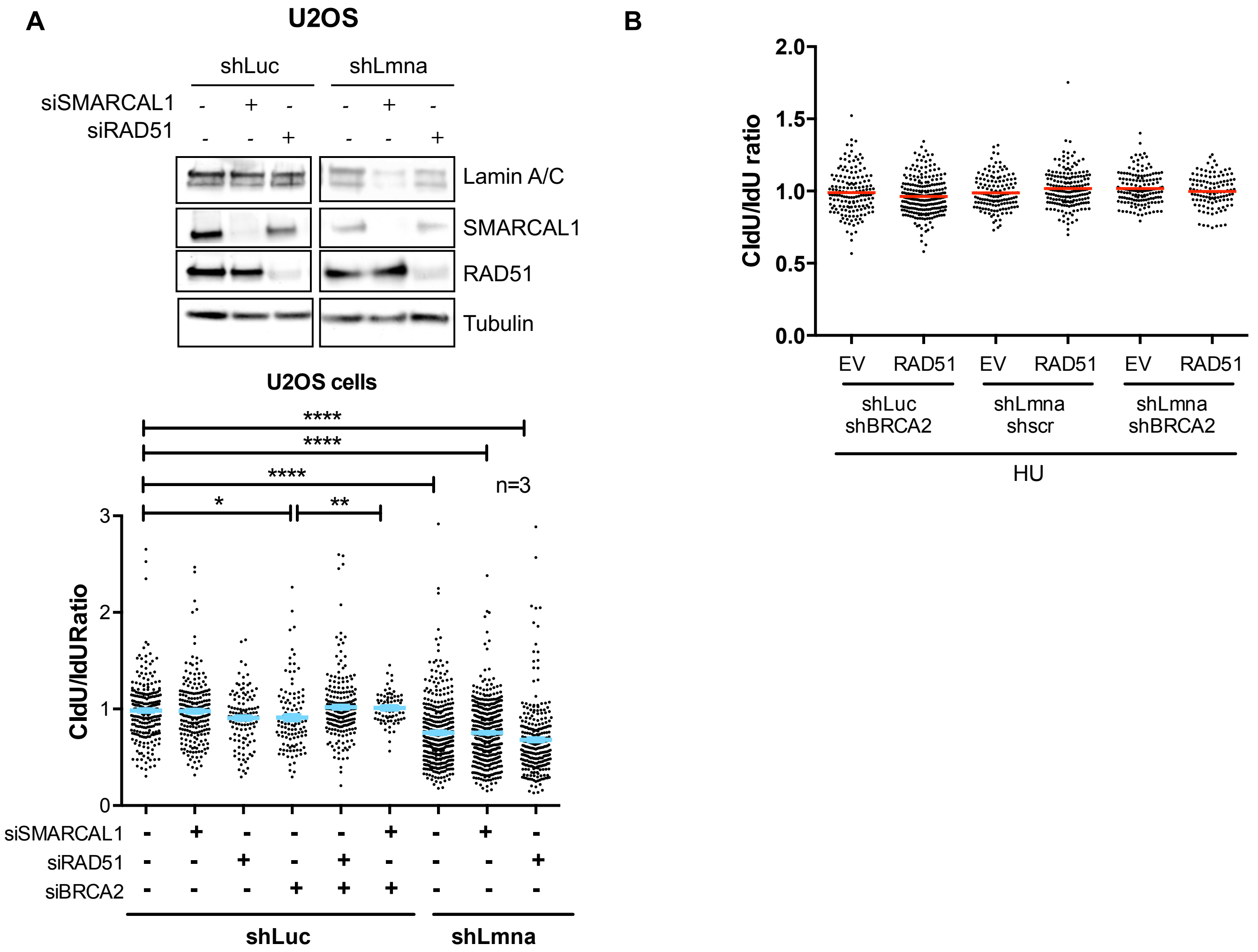


**Figure S5, related to Figure 3. SMARCAL1 neither RAD51 depletion rescue replication stress in lamin depleted cells. (A)** Immunoblots showing the transient depletion of SMARCAL1 and RAD51 via siRNAs in UO2S cells proficient (shLuc) or deficient for lamins (shLmna). Graph shows the result of DNA fibers (CldU/IdU ratio) performed in the different cells generated in (A). Note the RFI in BRCA2- and lamins-deficient cells and how depletion of SMARCAL1 or RAD51 only rescues in BRCA2 depleted cells. **(B)**Graph shows tract length ratio CldU/IdU in HEK293T cells depleted of lamins, BRCA2, or both combined, and overexpressing RAD51 or an empty vector control. Cells were labeled with IdU 20 min + CldU 20 min and not exposed to any drug treatment. Note how the ratio CldU/IdU is ~1 in all the cell lines and thus not affected by RAD51 overexpression. Data from three biological repeats and ~200 fibers measured in each experiment.

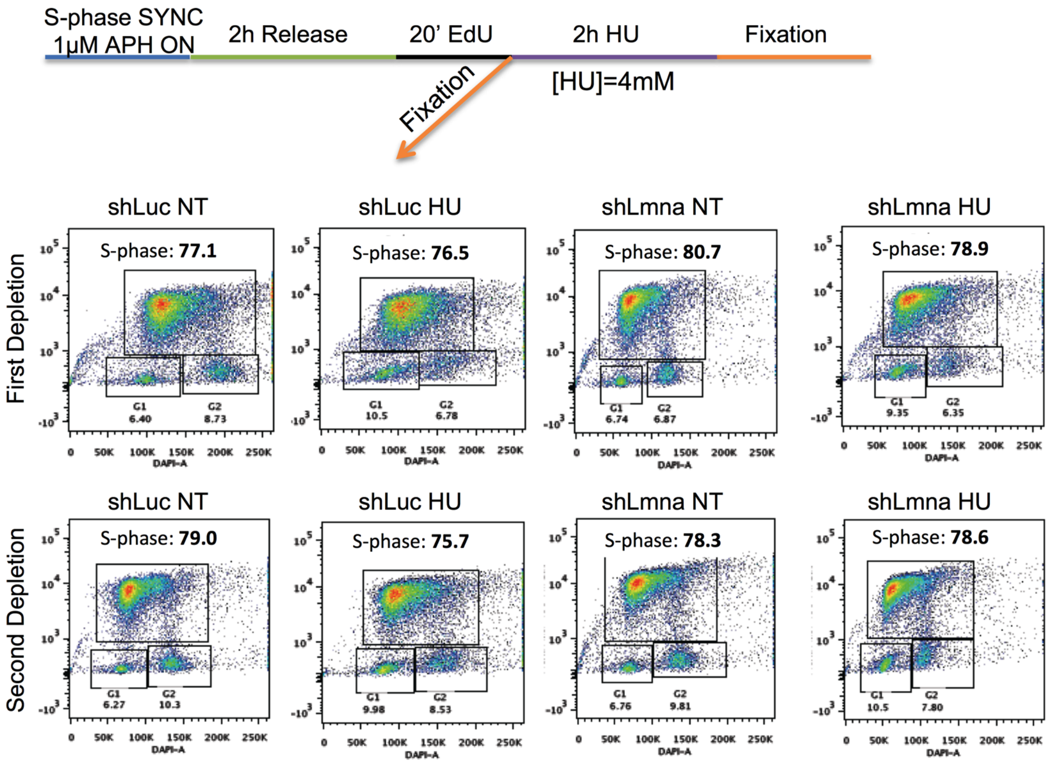


**Figure S6, related to Figure 5. Cell cycle profile of lamins-proficient and -deficient HEK293T cells synchronized in S-phase.** HEK293T cells depleted of lamins (shLmna) and control (shLuc) were treated with aphidicolin (APH) overnight to induce arrest in S-phase, followed by a release in complete media for 2 hours. Cells were then labeled with EdU for 20 minutes and either fixed immediately or after a 2-hour treatment with HU. Cells were analyzed for fluorescent DNA content and EdU content using the BD Biosciences LSR II flow cytometer, and the cell cycle profiles were created by the program FlowJo®. Note the efficiency of S-phase synchronization by this protocol, which is similar in the different conditions. Importantly, the fact that under these experimental settings cells incorporate EdU indicate that overnight exposure to APH, at the dosage used, does not damage replication forks as most cells are able to resume DNA replication. These measurements are essential to exclude the possibility that the phenotype (lack of recruitment of protective/remodeling factors) observed in lamin-deficient cells is due to the synchronization process per se rather than to the induced replication stress (HU in this case).

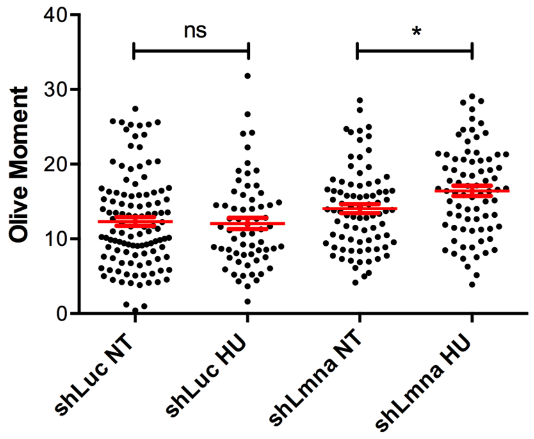


**Figure S7, related to Figure 6. DNA damage caused during HU treatment of lamins-proficient and -deficient cells.** Lamins-depleted and control HEK293T cells were subjected to the treatments used in the iPOND analysis (4mM HU for 2 hours) or no treatment (NT) as control. Cells were processed for Neutral Comet Assay to monitor the amount of DNA DSBs generated in response to RS. Graph shows average ± SEM of DNA DSBs in 3 biological repeats.
